# Structural and evolutionary determinants of Argonaute function

**DOI:** 10.1101/2024.10.10.617571

**Authors:** Arndt Wallmann, Mathew Van de Pette

## Abstract

Members of the Argonaute protein superfamily adopt functions ranging from host-defense to mediating elaborate and multi-component post-transcriptional and epigenetic systems of control. Despite this diversity of biological roles, the Argonaute structural fold is highly conserved throughout all domains of life. This raises questions about how Argonautes evolved to adapt to this increasing complexity of function, while conserving features that are broadly shared across the phylogenetic tree.

Integrating structural, sequence, phylogenetic data and disease-related mutational data, we compiled a comprehensive study of the Argonaute evolutionary trajectory. By comparing Argonaute proteins across a diverse set of lineages and extensive evolutionary timescale, we identified universal and clade-specific sequence signatures and intra-protein contact networks that underlie the Argonaute structural fold, nucleic acid interface and protein-protein binding sites. We analyze how these features are affected by disease-related mutations and are fundamentally altered in the case of the Argonaute-like Med13 protein. With this work we gain better insights into how Argonaute function diversified in eAgos by redrawing the emergence of conserved molecular features that are associated with new biological functions.

## Introduction

Across the phylogenetic tree, Argonaute proteins play a diverse set of roles that range from host-defense to complex epigenetic regulation ^1–3^. A key feature of Argonaute function is its ability to bind small oligonucleotide guides to target complementary nucleotide molecules, making it an adaptable and specific protein module to mediate sequence-guided function in the cell. Importantly, target-binding can result in many distinct biological consequences depending on the host species, Argonaute clade or ortholog, guide-target sequence complementarity, or the involvement of additional molecular factors. As part of the prokaryotic immune defense, many prokaryotic Argonautes (pAgos) target mobile DNA from viruses or plasmids via DNA interference (DNAi) and thus appear to be specific for DNA-guides and targets ^4–7^. Other pAgos specifically use RNA guides to recognize DNA targets and vice versa ^8–11^. Beyond prokaryotic immunity, pAgos have also been functionally linked to DNA replication and repair ^12–14^. Due to the high diversity of the pAgo family, the full scale and breadth of their functional and biological capacity remains largely unexplored. While 17% of eubacterial and 25% of archaeal genomes contain *ago* genes, almost all eukaryotes have Argonaute proteins ^15^. The broad functional repertoire of eukaryotic Argonautes (eAgos) includes RNA-guided RNA interference (RNAi), guiding DNA and histone methylation and targeting of viral RNA genomes ^16–18^.

Despite having diverged significantly in their sequence, eAgos and pAgos adopt a conserved bilobed fold with an N-terminal lobe (N-lobe) containing an N, a PIWI-Argonaute-Zwille (PAZ) and two linker (L1, L2) domains and a C-terminal lobe (C-lobe) formed by a Middle (MID) and a P-element Induced Wimpy Testis (PIWI) domain (Fig. 1a)^19,20^. The PAZ and MID domains bind the 3’ and 5’ ends of the guide oligonucleotide, respectively, and the PIWI domain contains the nuclease active site with a catalytic tetrad (DEDX, where X can be H/D) that typically cleaves the target between the 10th and 11th guide nucleotide ^21^. In most studied Argonautes, MID and PIWI domains pre-order the first several nucleotides on the guide, called the ‘seed’ region, in a helical conformation, which decreases the entropic cost for target binding ^22,23^. Duplex formation between the guide and target is often accompanied by opening of the Argonaute fold where the PAZ and parts of the N domain move as a discrete rigid body relative to the MID and PIWI domains ^4,24^.

**Fig. 1.**
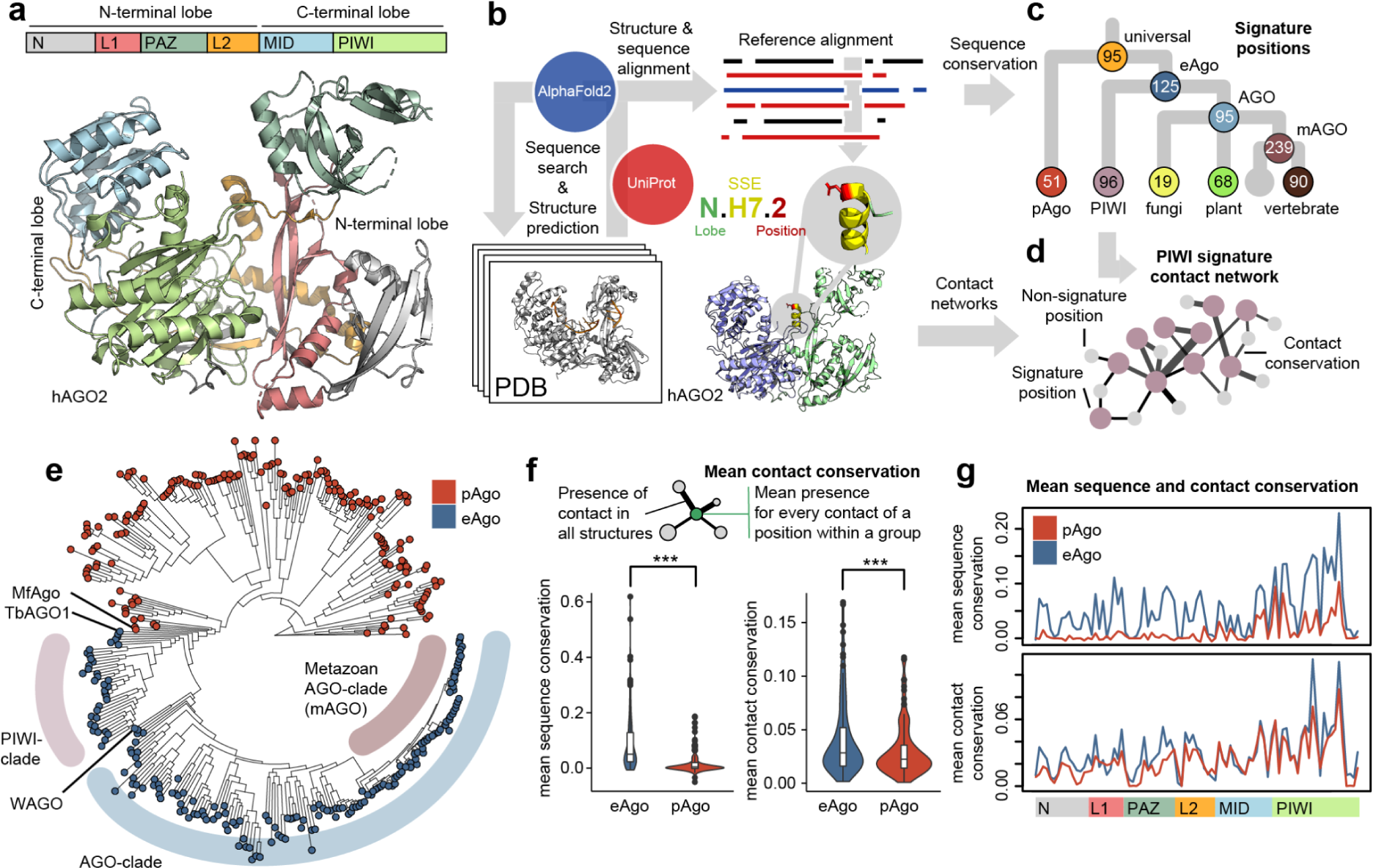
Overview of the Argonaute structural architecture, methodological approach and phylogeny. **a** Domain architecture of the long prokaryotic and eukaryotic superfamily of Argonautes. The bilobal structure consists of an N-terminal lobe containing an N, L1, PAZ and L2 domain and an C-terminal lobe with the MID and PIWI domain. The example structure shown is human Argonaute 2 (hAGO2, PDB: 4OLB). **b** Schematic overview of approach taken to create reference alignment (see Methods section). The Argonaute topology naming system identifies each position in the alignment by lobe, secondary structure element and position. **c** Using the alignment, the emerging conserved positions (signature residues) were determined for selected groups across the Argonaute phylogenetic tree. **d** Analyzing the local conserved structural environment of the signature positions produces a signature contact network for each group (e.g. PIWI-clade signature contact network). **e** Dendrogram of Argonaute based on the reference alignment with PIWI, AGO and metazoan AGO-clade highlighted (see Supplementary Figure 2 for a fully annotated dendrogram). **f** Mean sequence and contact conservation across all positions and plotted across the Argonaute domains (**g**).

Prokaryotic Argonautes can be phylogenetically divided into long-A and long-B pAgos and short pAgos, which only contain MID and PIWI domains ^15^. Sequence analysis indicates that long-B pAgos and short pAgos both have lost the ability to cleave targets, while long-A pAgos display an intact catalytic tetrad ^15,25^. eAgos include Trypanosoma and Nematode-specific Argonaute sub-families as well as the AGO and PIWI clade. While the AGO-clade proteins utilize micro RNA (miRNA) and short-interfering RNA (siRNA) to regulate mRNA expression, the PIWI-clade uses PIWI-interacting RNA (piRNAs) to protect the germline from transposable elements ^26,27^. In animal AGO-clade proteins, seed complementarity (nt 2–8) is sufficient for target recognition, whilst supplementary targeting (nt 12–17) has only a weak influence on target affinity ^24,28^. Plant AGOs use longer recognition elements (> 13 nt) ^29,30^. Extensive target complementarity to the 3’ end paired with central mismatches induces target-directed miRNA degradation (TDMD), which regulates miRNA stability in animals ^31^.

PIWI-clade Argonautes are required for fertility in many studied species ^26^. While the clade appears to have been lost in plants and fungi, some unicellular eukaryotes like amoebas and ciliates possess PIWI-clade proteins, where they instead appear to fulfill AGO-clade-like functions ^32,33^. In metazoa, PIWI-clade canonical seed pairing was recently shown to be dispensable for target recognition and PIWI-directed slicing tolerates mismatches to any nucleotide of the guide requiring a minimum of 15-nucleotide contiguous complementarity ^34^. Structurally, PIWI-clade proteins were shown to have a distinct interface between the L1, L2 and PAZ domains while possessing a widened central cleft region ^35–37^.

Based on sequence analysis of its PIWI module, the Mediator complex Med13 protein was proposed as a third divergent clade in the eukaryotic PIWI/AGO superfamily ^38^. Med13 is part of the Cdk8 kinase module (CKM), which is a dissociable subcomplex of the Core Mediator complex that can sterically block interaction with RNA polymerase II ^39,40^. Recent work on the *Saccharomyces cerevisiae* CKM complex revealed remarkable structural similarities between the Med13 and Argonaute domain architecture ^41^. However, it is still unclear if Med13 possesses fundamental eAgo functions like small nucleic acid binding or target matching.

Previously, structural and sequential differences and similarities between Argonautes have been noted in pairwise comparisons, but an integrated and systematic analysis of the available structure and sequence data across the phylogenetic tree has been lacking ^9,35,41^. Although having diverged substantially on the sequence level, the Argonaute structure is restrained by its function to bind and match small oligonucleotide guides to complementary targets. Hence, a systematic structure-guided comparison promises to yield better outcomes than a purely sequence-based analysis. Moreover, the recent development of novel machine learning-based structure prediction tools like AlphaFold2 (AF2) has enabled new approaches for evolutionary in-depth protein contact analysis ^42^. In the analysis described here, we identified and analyzed long Argonautes across the Argonaute/PIWI superfamily and pinpointed signature residues that emerge at specific points in the evolutionary trajectory of Argonautes. We determine the signature residues and their conserved contact network from all long Argonautes down to vertebrate AGO-clade Argonautes and integrate this data with structural data on nucleic acid and protein-protein interfaces. Our analysis determines critical positions of the guide seed binding region and emerging eAgo-specific features of the second half of the guide interface. In addition, we identify distinct molecular similarities between the PIWI-clade and pAgos. Using cancer and large-scale population variation data, we cross-reference and assess disease-related mutations in a structural and phylogenetic context and show that many target the nucleic acid interface. Finally, we apply our analysis on the Argonaute-like Med13 protein, uncovering how many of the universal and eAgo-specific signatures of the nucleic acid interface have diverged substantially in this protein family.

## Results

### Integrating sequence and structural data via a structure-guided Argonaute alignment

To enable meaningful comparison of the structurally equivalent residues between Argonautes from evolutionary distant organisms, we created a comprehensive structure-guided sequence alignment of long pAgos and eAgos. This reference alignment was created with an iterative approach using the available protein structures from the Protein Data Bank (PDB; Supplementary Data 1), profile-based sequences searches, and structural alignment of high-quality AF2 models (Fig. 1b, see Methods). Importantly, our structure-based search strategy enabled us to identify and include highly diverged Argonaute sequences that may have been missed through purely sequence-based alignment (total of 17,461 sequences, see supporting data). We find that the vast majority of sequences are of eukaryotic origin, and that the average pAgo protein size is lower than that of eAgos (Supplementary Fig. 1a,b). Interestingly, we find 35 protein sequences with multiple consecutive long Argonaute modules in plants, animals and Amoebozoa (Supplementary Data 2). While for most pAgos we only find one Argonaute per species, there is a trend of gradual expansion in plants and animals with multiple Argonaute sequences per species (Supplementary Fig. 1c). Utilizing the Orthologous MAtrix (OMA) database, we find a similar emergence of Argonaute paralogs in eukaryotes, which highlights the diversification of functions in plants and animals (Supplementary Fig. 1d). In agreement with previous studies, we find many nematode species with more than 20 Argonaute proteins (e.g. *Caenorhabditis elegans, Panagrellus redivivus*), but also some plant species (e.g. *Brassica napus, Glycine soja*) and selected fungi species (e.g *Diversispora epigaea*, *Glomus cerebriforme*) display similarly high Ago sequence numbers. The sequences included in the final reference alignment were selected based on sequence similarity and phylogenetic considerations to ensure that the alignment represents the broad variability of long Argonautes from a diverse set of organisms (Supplementary Fig. 1e). The reference alignment contained a total of 366 Argonaute sequences with a 1:1 ratio of eAgos and pAgos (183 Eukaryota, 37 Archaea, 146 Bacteria, Supplementary Data 2).

To enable comparability between positions in the alignment, we devised an Argonaute residue naming scheme, based on the overall protein topology and secondary structure of human Argonautes (hAGOs), in analogy to similar approaches studying G protein-coupled receptors and TATA-box binding protein ^43,44^. The naming scheme includes the lobe of the position, the number of the respective secondary structure element (SSE, H: helix, S: strand, L: loop) and the residue position within that SSE (e.g. N.H7.2 for the second position of the 7th helix in the N-lobe; Fig. 1b; Supplementary Fig. 1f). For all Argonautes lacking an experimental protein structure, we computed a structural model using AF2 (see supporting data). We find that the predicted Argonaute structures have a high overall prediction score with a mean pLDDT confidence score of 90 (Supplementary Fig. 1g). Indeed, in the case of the *Pseudooceanicola lipolyticus* Argonaute (PlAgo), AF2 produced a model highly similar to the experimental structure, which was published after the generation of the AF2 model (all atom RMSD of 2.16 Å, Supplementary Fig. 1h). To create an integrated contact network, the non-covalent residue contacts for each of the experimental protein structures and the AF2 models were calculated and referenced via the Argonaute topology (Supplementary Fig. 1i). For each phylogenetic subgroup we then defined distinctly conserved residues in that group (signature positions, Fig. 1c, Supplementary Data 3) and the conserved contacts this set of residues is involved in (signature contact network, SCN; Fig. 1d, Supplementary Data 4) to identify emerging and shared molecular features (see Methods). To minimize errors and inaccuracies known to arise from AlphaFold2 predictions, we applied a rigorous confidence threshold to the analysis of the models, summarized atomic interactions into residue-residue interactions and only used groups of models for analysis ^45^.

### Argonaute phylogeny and the relationship between sequence and contact conservation

We created a dendrogram based on the structure-guided reference alignment to structure the data along phylogenetic relations. The resulting tree shows a strict separation between prokaryotic and eukaryotic Argonaute sequences (Fig. 1e, fully annotated tree in Supplementary Fig. 2). Archaeal Argonautes and bacterial sub-families are scattered over the pAgo branch. This is indicative of horizontal gene transfer as one of the driving mechanisms behind the spread of Argonautes in prokaryotes and is consistent with previous reports ^46^. The closest relative to eAgos is a pAgo from an extreme thermophile Euryarchaeota (*Methanotorris formicicus*, MfAgo). The eAgo family is divided into the PIWI-clade and the AGO-clade with the metazoan AGO-clade (mAGO) representing a highly conserved subgroup. The single *Trypanosoma brucei* (TbAGO1) representative included in the analysis is positioned as a monophyletic group closest to the pAgo-clade, which may reflect its early divergence from eukaryotes as a lineage-specific duplication. The single Nematoda-specific Argonaute (WAGO) clade representative is part of the AGO-clade branch but separate from the mAGO-clade as has been shown before ^46^.

We next analyzed how sequence and structure is conserved within the two main groups of our dataset, eAgos and pAgos. Sequence conservation is significantly higher in eAgos than in pAgos (Fig. 1f, bottom). As a measure of structural conservation we computed the mean contact conservation of all non-covalent contacts for each position within a selected group of structures and models (Fig. 1f top). Overall mean contact conservation is also higher in eAgo than pAgo structures (Fig. 1f, bottom) and positively correlates with sequence conservation in both (Supplementary Fig. 3a). However, when mapped across the whole protein, pAgo contact conservation follows eAgo contact conservation much more closely than sequence conservation (Fig. 1g). This finding illustrates how the overall structural fold remains largely preserved and how pAgos sample a larger sequence space than eAgos.

### Universal properties of the Argonaute fold

In order to identify key features of the Argonaute fold, we determined the universal signature positions and universally conserved contacts across the 366 Argonaute sequences. Using ancestral sequence reconstruction and sequence conservation scoring (see Methods), we identified 95 residues that are universally conserved across all Argonautes and are enriched in the conserved C-lobe (Supplementary Fig. 3b). We note that in our analysis the first three positions of the catalytic tetrad DEDX motif are classified universal, even though they diverged within the branch of long-B pAgos (Supplementary Fig. 3c). We next calculated conserved contact networks for eAgos and pAgos separately and integrated the common interaction network with the universally conserved residues, to determine the universal SCN (Fig. 2a). This universal SCN includes structural features that are widely conserved across all Argonautes such as the hydrophobic cores of the MID, PIWI and PAZ domain (Fig. 2b,c). Most universally conserved positions are deeply embedded in the Argonaute structural core and thus mediate a substantial amount of contacts in the structure (Supplementary Fig. 3d). Next, we used the available experimental structures of Argonautes to determine the nucleic acid (NA) interface residues (Supplementary Data 5). Argonaute NA interface data is often incomplete or ambiguous since targets and guides are not fully resolved in most structures (Supplementary Fig. 3e,f) or show structural or conformational variability (e.g. duplex formation, TDMD, supplementary targeting). Hence, we consider a position or contact to be shared if it is present in at least one pAgo and eAgo structure (Fig. 2d, Supplementary Fig. 3g,h). Of those 111 shared NA interface residues, 19 are universally conserved and make shared contacts to distinct guide or target positions (Fig. 2d,e). These positions are located in the 5’ MID and seed binding region, catalytic region and the PAZ domain (Fig. 2e, bottom panel). We identify two clusters of universally conserved positions that contact the guide seed: One is located in the MID domain contacting guide 1-3 and is responsible for 5’ guide end binding. The second contacts the end of the seed (guide 6-8) and guides it as part of the helical pre-ordering. Overall, our results identify the universal contacts that build the structural core and highlight how guide seed interactions represent a highly conserved feature across Argonautes.

**Fig. 2.**
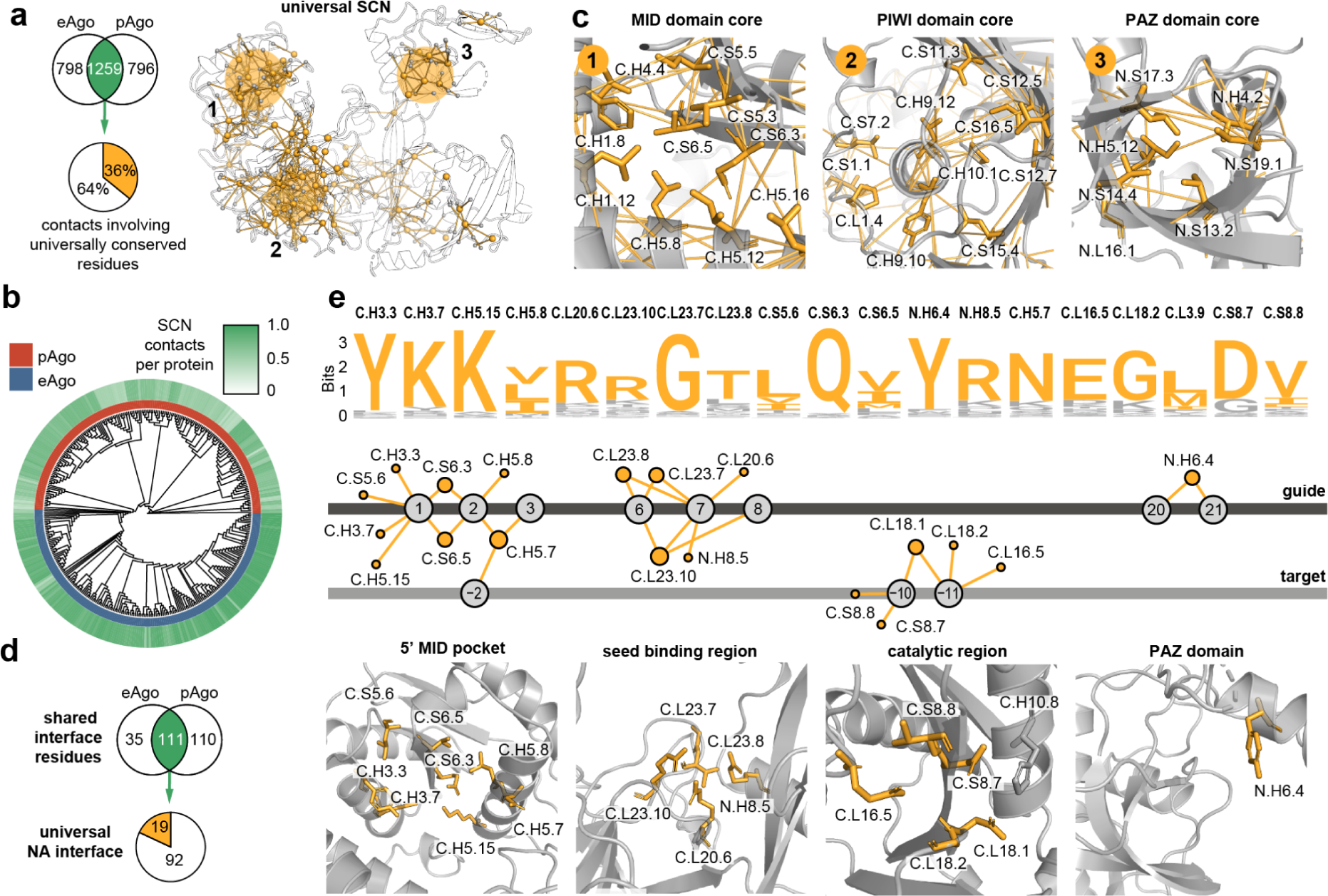
Universal structural features of Argonautes. **a** Venn diagram of overlap between conserved contacts in pAgos and in eAgos and pie diagram of the fraction of common contacts with at least one universally conserved residue. The resulting conserved contact network is projected onto the hAGO2 structure. **b** Heatmap showing the proportion of universal SCN contacts for each Argonaute across the phylogenetic tree. **c** Examples of positions in the universal conserved contact network as highlighted in **a**. **d** Guide and target interface of eAgos and pAgos derived from structural information in the PDB (see Methods). Overlap of shared interface residues between pAgos and eAgos (111) and proportion of universal residues in the shared interface with shared contacts (19). **e** The universally conserved NA interface is visualized using a logo-sequence plot (top), network representation (middle) and structure (bottom, PDB: 4OLB).

### Conserved molecular features of pAgos and eAgos

Next, we identified the signature positions that showed high conservation in either eAgos or pAgos, but were not conserved or were conserved as a different residue, in the corresponding group. This resulted in 125 positions for eAgos and 51 positions for pAgos with 18 positions shared but conserved as different residues (Fig. 3a). The signature residues were integrated with the respective contact network to yield the SCNs of pAgos and eAgos, respectively (Fig. 3b,c). The eAgo SCN covers both the C- and N-lobe, while most of the pAgo SCN is located in the C-lobe. We find several regions that display distinct local sequence and structural features in the core and on the surface of pAgos and eAgos (Supplementary Fig. 4a,b). We also find that in contrast to pAgos, eAgos carry N-terminal ends that are predicted to have a higher protein disorder content (probability value > 0.5, Fig. 3d). Consistent with this trend, pAgos have been shown to carry additional N-terminal domains that may modify nucleic acid binding, while the highly disordered eAgo termini may contain contact sites for eukaryotic-specific interaction partners ^15,47^.

**Fig. 3.**
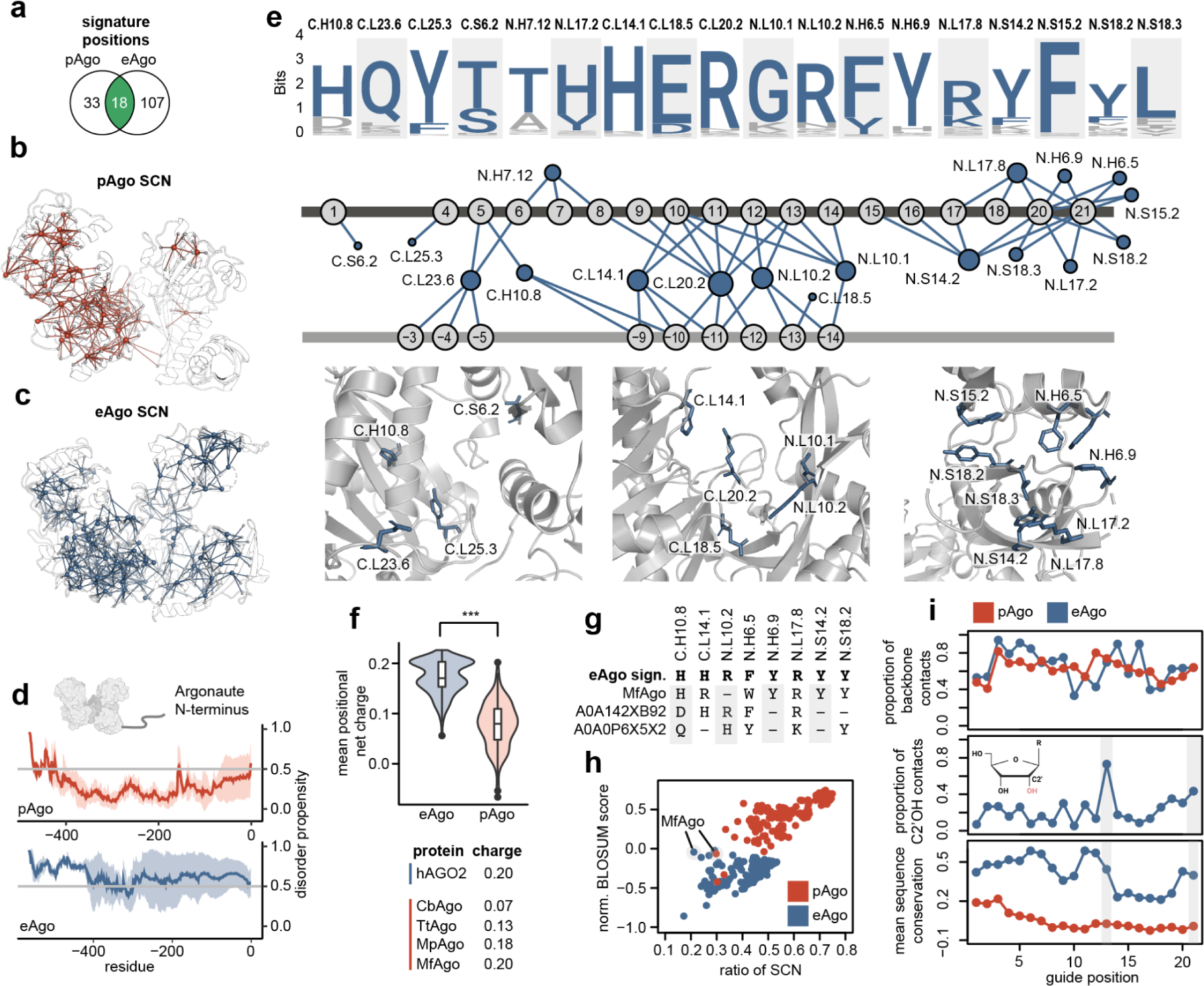
Molecular signatures in the nucleic-acid interface in pAgos and eAgos. **a** Signature positions overlap between pAgos and eAgos. **b,c** Structural overlay of the pAgo SCN (red, TtAgo, PDB: 3DLB) and eAgo SCN (blue, hAGO2, PDB: 4OLB). **d** Predicted Argonaute N-terminus disorder content for all pAgos and eAgos included in this study. **e** eAgo NA interface signatures illustrated in the logo-plot, NA contact network and labeled cartoon representation. **f** Violin plot of mean positional net charge for pAgo and eAgo shared NA interface positions. A selection of proteins is listed together with their interface charge value. **g** eAgo NA signature positions of MfAgo and two closely related pAgos. **h** Individual pAgo proteins plotted for normalized signature eAgo (blue) and pAgo (red) BLOSUM score and SCN ratio. MfAgo (UNIPROT: H1KXR6) is annotated in the plot. **i** Proportional ratio of all contacts to guide backbone resolved by guide position in eAgos and pAgos for all included structures (top). Proportional contacts of C2’OH contacts of all guide backbone contacts in eAgos (middle). Mean sequence conservation of NA interface resolved by guide position (bottom). The 13th position and the 3’end of the guide are highlighted.

Next, we integrated the NA interface data with eAgo and pAgo signature positions. Many pAgo proteins show unique NA interactions (Supplementary Fig. 4c) and the pAgo branch shares few pAgo-signature positions in the NA interface (Supplementary Fig. 4d). In contrast, we identified a total of 18 NA interface positions among the eAgo signature positions, which are mainly located near the endonuclease active site and the guide 3’ end (Fig. 3e). When assessing common NA binding positions, the average NA interface is significantly more positively charged in eAgos than in pAgos (Fig. 3f) and this difference emerges in the 3’ half of the guide (Supplementary Fig. 4e). Among pAgos, we find that the NA interface of MfAgo carries one of the highest overall charges and shares many of the eAgo-signature positions of the NA interface (Fig. 3f,g). Indeed, when scoring the consensus motif of the eAgo and pAgo signatures against all pAgo sequences, only MfAgo scores better for the eAgo than the pAgo signature supporting its phylogenetic position close to eAgos (Fig. 3h). Interestingly, the corresponding structural model contains only few (21%) of the eAgo SCN contacts, indicating a different local structural environment of the signature eAgo positions.

All studied eAgos use RNA guides for targeting ^48,49^ and hAGO2 has directly been shown to display RNA over DNA-guide preference ^50,51^. As guide binding is mostly sequence independent and most NA contacts are with the NA backbone (Fig. 3i, top), we reasoned that the ribose backbone C2’ OH in RNA guides may be of relevance to guide selection. Thus, we analyzed the RNA backbone contacts across the available eAgo structures. The 13th position and the 3’end of the guide display many direct contacts to the C2’ OH (Fig. 3i, middle) and show overall conservation of the involved residues (Fig. 3i, bottom). Many of those contacts are mediated by the eAgo signature positions N.L10.2 (guide positions 13), N.S18.2 and N.S15.2 (3’ end), which consequently represent candidate positions implicated in guide selection. Taken together, the conserved changes to the eAgo NA interface are mainly found in the second half of the guide interface, underlie a positive charge in that region and may control RNA guide selection.

### The diverging structural signatures of the PIWI- and AGO-clade

Multiple structural studies have highlighted unique features of the PIWI and AGO-clade family ^35–37,52^. We identified 96 and 95 signature residues that are unique to the PIWI or AGO-clades, respectively, and determined the SCNs by analyzing their conserved contact environment (Fig. 4a,b). The N.L23 loop forms the central gate in AGOs and inhibits target-guide interaction in the central regions of the miRNA. In contrast, the central cleft in PIWIs is widened through the formation of a helix in N.L23, which allows for more pairing ^37^. We do not find PIWI signature positions within this region, but the alpha-helical backbone contact from N.H7.1 is part of the PIWI SCN (Fig. 4a,b, panel 1). Conversely, N.L23.10 is conserved as a leucine in the AGO-clade which may indicate that its contacts with N.H7.5 stabilizes the central gate. The PIWI signature residues C.L23.4 and C.L23.9 are more bulky in PIWI than in AGO and thus cannot kink to cradle the guide at positions 5 and 6, leaving that part of the seed not pre-organized in PIWI (Fig. 4a,b, panel 2) ^37^.

**Fig. 4.**
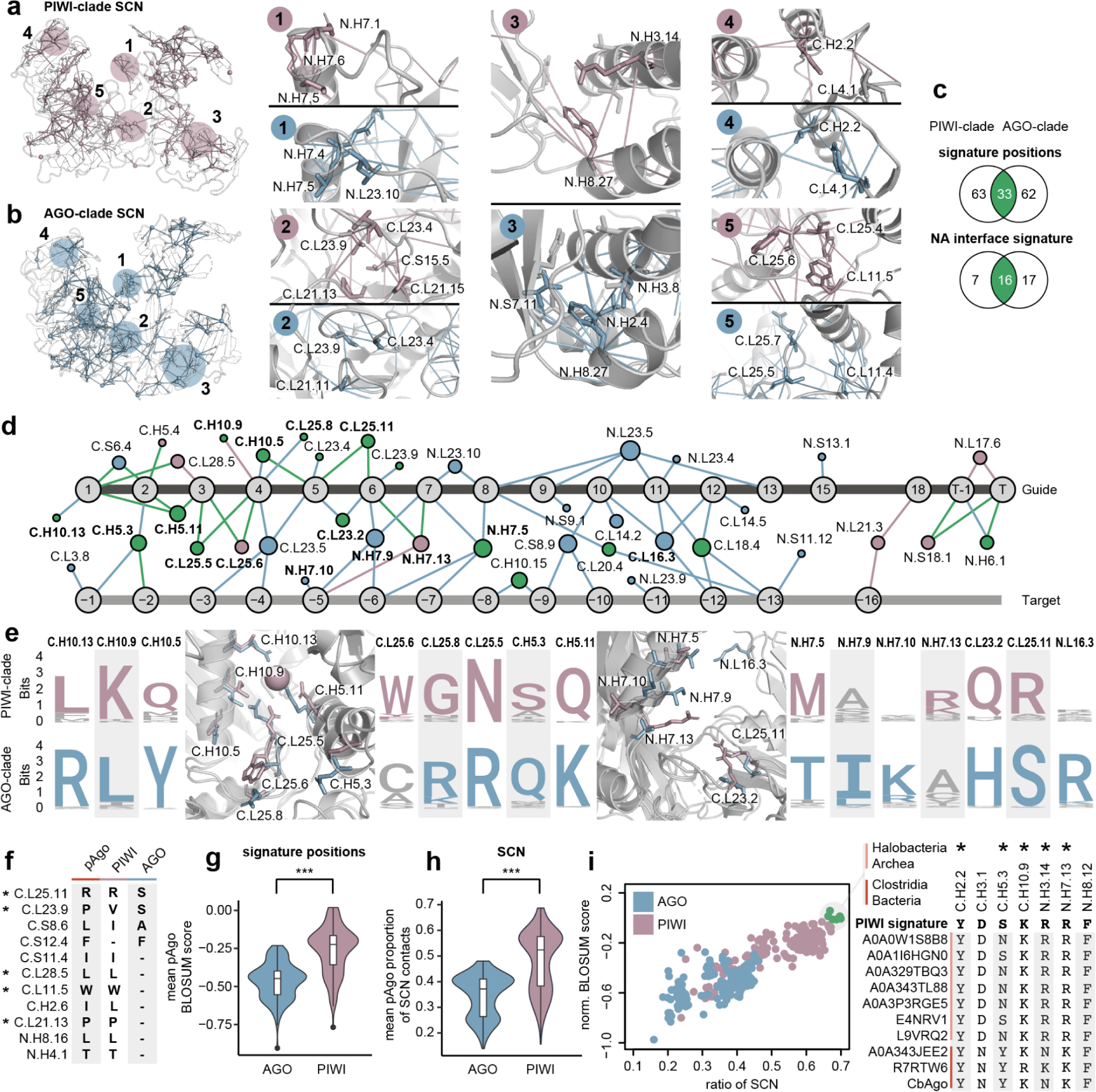
PIWI and AGO signatures, NA-binding and comparison with pAgo. **a** Structural overlay of the PIWI-clade SCN with numbered and highlighted regions in pink. **b** Structural overlay of the AGO-clade SCN with numbered and highlighted regions in blue. **c** Total signature position overlap and respective NA interface signature overlap between the AGO and PIWI-clade. **d** PIWI-and AGO clade signature NA interface depicted as a contact network with shared (green), and PIWI (pink) or AGO-specific (blue) contacts. **e** Logo plot and cartoon representation of selected positions in the PIWI/AGO NA interface. **f** Comparison between the pAgo, PIWI and AGO signature positions and their respective consensus residue. Positions marked with a star are shown in **a**, **b** or **e**. A dash represents a missing signature at the respective position. **g** Normalized BLOSUM scoring of the AGO and PIWI signature against all pAgos from this study. **h** Ratio of detected SCN contacts by pAgos for the PIWI and AGO SCN. **i** Individual pAgo proteins mapped for normalized signature BLOSUM score and SCN ratio for PIWI and AGO. Highest scoring group of ten pAgos for PIWI contains members from Halobacteria and Clostridia (highlighted green) and shows sequence similarity to PIWI-signature positions. Positions marked with a star are shown in **a**, **b** or **e**.

Previously, the N-L1-L2 interface had been highlighted as distinct between the AGO and PIWI-clade, leading to a more open N-conformation in PIWI ^35^. We find that AGO-clade signature positions N.S7.11, N.H2.4 and N.H3.8 form a conserved extended hydrophobic interface and support the more compact conformation. In contrast, PIWIs harbor a signature position, N.H3.14, carrying a positively charged lysine or arginine residue that may weaken the hydrophobic interface and thus stabilize the more open N domain arrangement (Fig. 4a,b, panel 3).

We also find major differences that support the distinct arrangement of the PAZ domain in PIWI and AGO (Supplementary Fig. 5a-c), which may in part reflect the distinct lengths of their guide RNAs. In addition, we detect minor but highly conserved differences that do not seem to affect the overall structure, but may give clues to the phylogenetic origin of the two clades (Fig. 4a,b panel 4,5).

Of all the respective signature positions, 33 positions overlap between the PIWI and AGO-clades and are conserved as different residues (Fig. 4c). Based on the available structures (Supplementary Fig. 5d), almost half (16) of these positions are located in the NA interface and thus, can directly impact NA binding (Fig. 4c, Supplementary Fig. 5e). Hence, we mapped out the respective position-specific guide and target interactions of the PIWI and AGO-clade signature interfaces (Fig. 4d,e). Many of these diverging signature positions and the resulting clade-specific contacts can be found among the seed-interacting residues of the C.L25/C.L23 loops and the N.H7 helix. In contrast, the NA positions 8-17 are exclusively contacted by AGO-signature interactions, which include positions from the above-mentioned N.L23 that forms part of the central gate (N.L23.4/5/9/10), but also the C.L16 (PIWI-loop) and C.L14 loop. While the diverging signature positions C.H10.13, C.H10.9, and C.H5.11 have been linked to metal-dependent 5’ guide recognition in PIWI, the AGO-clade mediates these interactions through the charged residues C.H10.13 and C.H5.11 ^36^.

As metal-dependent 5’ guide recognition and other features are shared between the PIWI-clade and some pAgos, it was proposed that PIWI represents an ancient variant of eAgos ^2^. Hence, we investigated how PIWI- and AGO-clade sequence signatures and SCNs compare to the pAgo group. We find that the signature positions of pAgos and PIWIs contain many identical or similarly conserved positions (Fig. 4f). Among those is C.L11.5, which is part of a group of polar residues in AGOs that recognize exchangeable internal water molecules that form the so-called loop-associated key estuary 1 (LAKE1) ^53^. PIWI has been shown to possess conserved hydrophobic residues in these positions. We find that this seems to be a shared feature with pAgos, which also predominantly display a tightly packed hydrophobic core in that region (Supplementary Fig. 5f). This suggests that the domain convergence upon guide loading described in hAGOs is likely a specific property of the AGO-clade and highlights the structural similarity between pAgos and PIWIs.

When scoring the PIWI-and AGO signature positions and their contacts against pAgos, we find that both are significantly higher for PIWI (Fig. 4g,h). Interestingly, a group of pAgos from Halobacteria archaea and Clostridia bacteria display the highest similarity to PIWI (Fig. 4i).

Collectively, our analysis finds distinct differences in the seed interface between the PIWI- and AGO-clade and suggests that, compared to the AGO-clade, the PIWI-clade shares more molecular features with pAgos.

### The GW-protein interface emerges in the AGO-clade

In animals, GW182 proteins recruit AGO-clade Argonautes into P-bodies via an intrinsically disordered Argonaute binding domain (ABD) harboring multiple WG/GW repeats ^54–56^. As GW-repeat proteins have been reported to interact with Argonautes from plants, fungi and even the PIWI-clade, the interaction was proposed to be widely conserved among Argonautes (Fig. 5a) ^57–61^. Three evenly spaced tryptophan (TRP1-3) binding pockets in the hAGO2 PIWI domain mediate stable interactions with the human GW182-protein TNRC6B (Fig. 5b) ^62,63^.

**Fig. 5.**
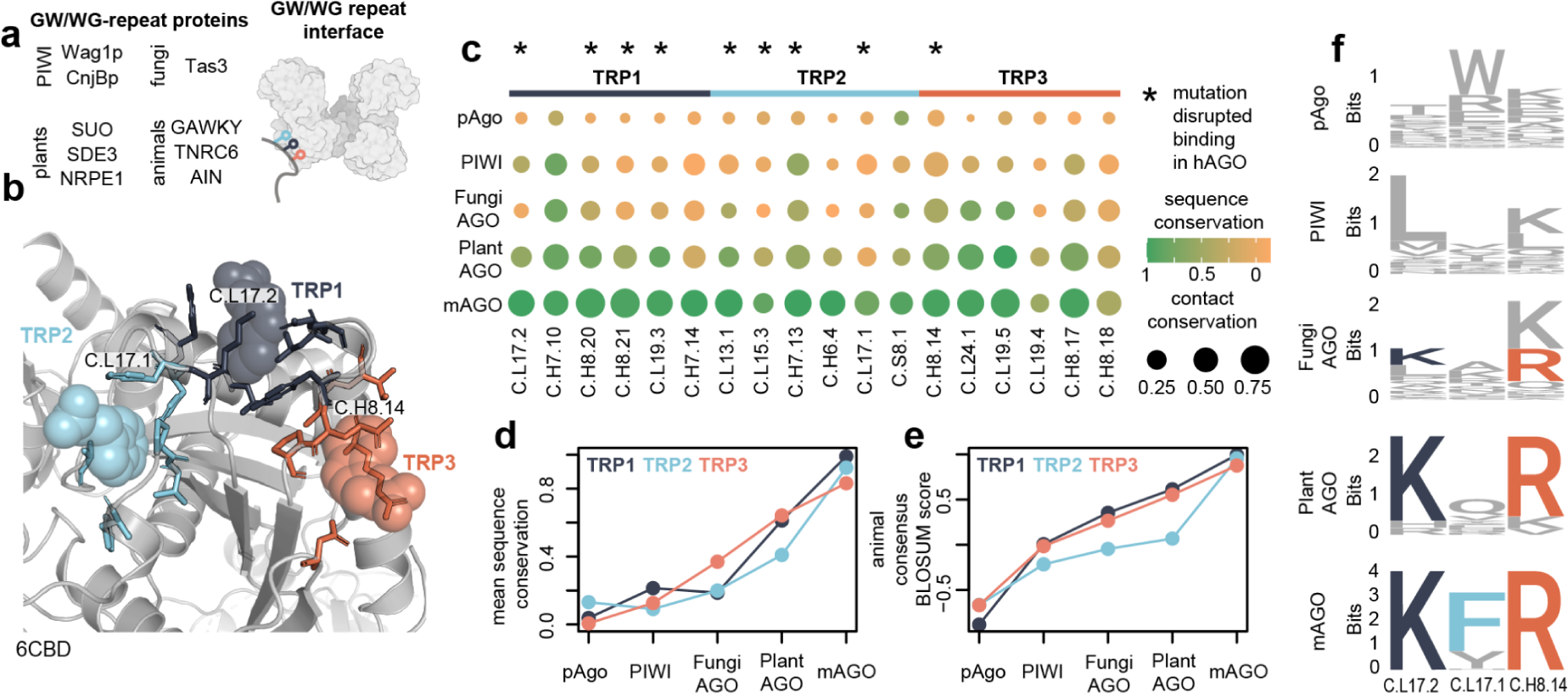
Signatures of the PIWI and AGO-clade and the GW-protein interface. **a** Candidate proteins proposed or shown to interact with eAgos via GW/WG-repeat modules **b** The interface location in the PIWI domain (hAGO2, PDB:6CBD). **c** Dotplot of interacting residues for the three tryptophan binding sites showing sequence (color-coded) and contact (size) conservation. Positions marked with a star interfered with GW-binding when mutated in previous studies. **d** Mean sequence conservation for all three tryptophan binding sites for pAgos and eAgo subgroups. **e** Mean normalized similarity score for the consensus mAGO sequence of all three tryptophan binding sites for pAgos and eAgo subgroups. **f** Logo plots of three key sites for each of the tryptophan binding pockets (Colour coded residues are annotated in **b**).

To study the emergence of the interface, we first identified the positions directly involved in interactions between the GW-protein TRPs and the three TRP binding pockets in hAGO2. Next, we analyzed the contact and sequence conservation of these positions (Fig. 5c, Supplementary Fig. 6a). All three binding sites are strongly conserved in mAGOs, while displaying high variability in pAgos and the PIWI-clade. When calculating the mean sequence conservation for each TRP binding pocket, we find a non-uniform increase in conservation across the AGO-clade (Fig. 5d). While the TRP3 pocket shows distinct conservation in fungi, the TRP2 pocket only matches its level of conservation in mAGOs. Scoring the mAGO consensus sequence of the three pockets against all Argonautes, we find that the mean similarity score of pocket TRP2 remains lower than the other two pockets across fungi and plants (Fig. 5e, Supplementary Fig. 6b). Indeed, in the available structure of Arabidopsis thaliana AGO10 a tryptophan at position C.L15.2.1 (732) occupies the TRP2 pocket, likely blocking the binding site, that is highly conserved in and specific to plants (Supplementary Fig. 6c). Figure 5f shows the logo plots of three positions that have each been shown to be essential for their respective pockets, when mutated in hAGO2 ^62,63^.

Collectively, the data suggests that despite reports of GW-binding Argonaute in PIWI-clade and fungi AGO-clade Argonautes, the binding sites remain remarkably variable in these groups and do not appear to belong to the shared functional repertoire of eAgos. Notwithstanding knowledge bias towards mAGO-GW interactions and the possibility of currently unknown GW-interaction sites in Argonautes, our analysis supports a model, in which GW-binding sites accumulated in eAgos and where three fully conserved TRP-binding sites are only a conserved feature of mAGOs.

### Argonaute missense mutations in disease

We conducted a literature and data review to evaluate the validity of our identified signature positions and to contextualize the positions with structural and phylogenetic information. We first evaluated 12 hAGO2 germline mutations associated with the Lessel-Kreienkamp (LESKRES) syndrome, which severely affects neurological development leading to delayed motor development and intellectual disability in patients ^64^. The mutations were reported to mainly impair RNA-induced silencing complex (RISC) formation or disrupt proper NA-interaction. All of the 12 mutated positions found in LESKRES syndrome are part of conserved Argonaute signatures (Fig. 6a). The positions C.L23.9 (760) and N.H7.8 (364) make contact with the guide and are part of the eAgo- and AGO specific NA interface, respectively. Thus, mutation of those residues likely affects NA-binding directly. While most positions are only conserved in eAgo and its subgroups, two of the analyzed mutations belong to the universal signature of Argonautes: C.L21.16 (G733R) and C.L11.2 (G573S). These two glycines form heavily conserved contacts across all Argonautes in the universal SCN (Supplementary Fig. 7a). Strikingly, C.L21.16 was the only characterized mutation that appeared to exhibit a near complete loss-of-function in most of the functional assays performed, including mRNA binding and P-body localization ^64^.

**Fig. 6.**
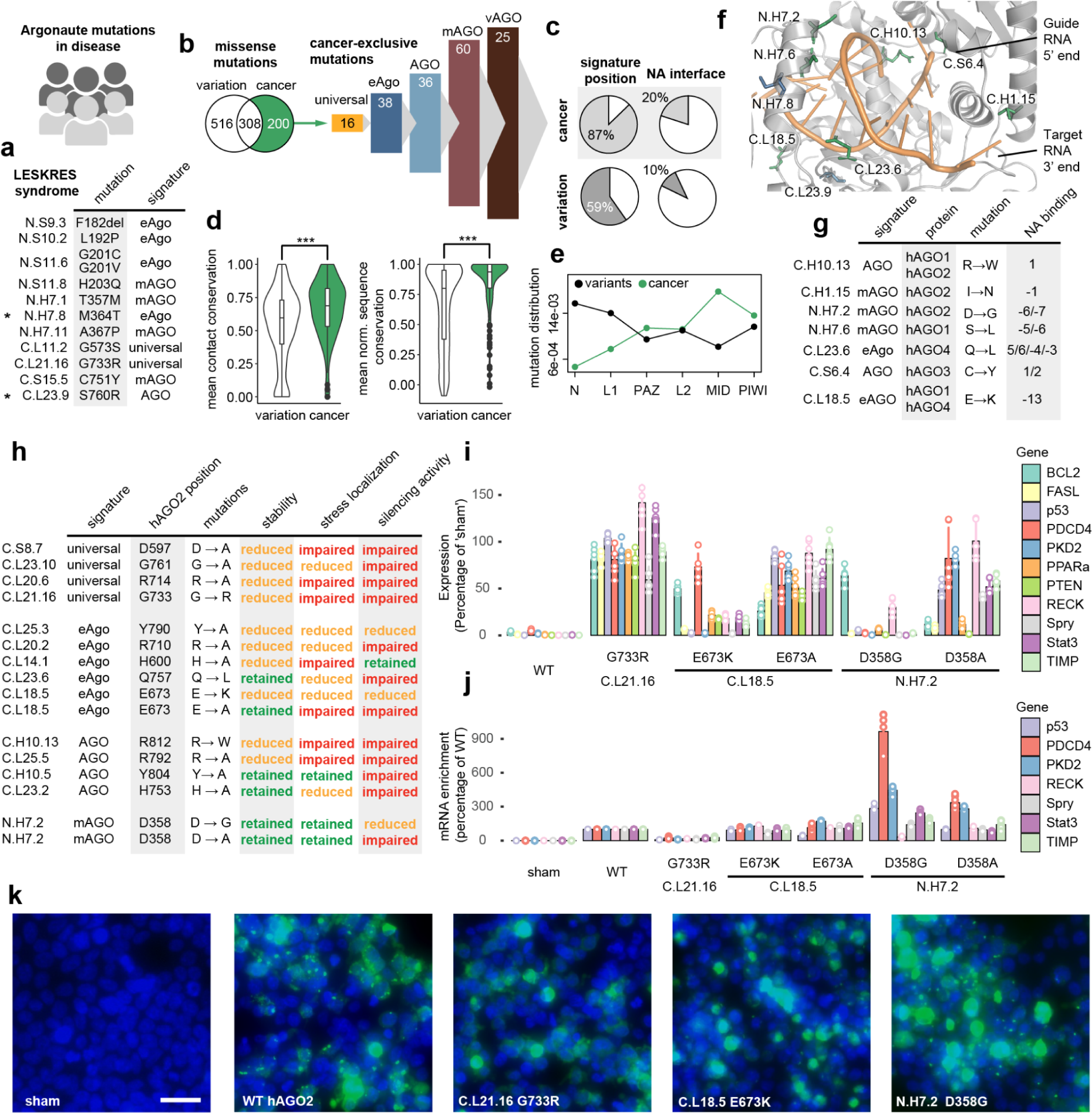
Argonaute missense mutations in disease. **a** List of mutations from LESKRES syndrome and their annotation to the Argonaute topology and signatures. **b** Venn diagram of missense mutation positions from population variation (gnomeAD) and somatic cancer mutation data (cBioPortal). 200 positions that are exclusive to the cancer mutational data (green) are assigned to the respective signatures (vertebrate AGO-clade: vAGO) **c** Proportions of cancer-exclusive mutations and population variation mutations (white in **e**) that affect signature positions or the nucleic acid interface. **d** Mean contact and sequence conservation of population variation and cancer mutations. **e** Distribution of the variation and cancer mutations across the Argonaute domains. **f** Examples of cancer mutations in the nucleic acid interface. The two mutations directly affecting guide-contacts from LESKRES syndrome (**a)** are colored blue in the structure, cancer mutations (**g**) in green. **h** Summary table for mutations hAGO2 used in functional assays. **i** RT-QPCR analysis of relative miR-21 target genes abundance following mutant hAGO2/miR-21 co-transfection. **j** RT-QPCR analysis of miR-21 target gene enrichment following mutant hAGO2/miR-21 co-transfection and subsequent hAGO2 immunoprecipitation. **k** Live cell imaging of hAGO2-GFP (green) containing stress granule (small speckles) formation following Arsenite exposure. Nuclear signal (blue) overlain. Scale bar 20µm.

In contrast to LESKRES syndrome, where mutations instigate a loss of function, growing evidence hints at an oncogenic potential of Argonaute proteins in cancer ^65^. We assessed hAGO somatic cancer mutation data sourced from the cBioPortal database and we referenced it to natural variation data from gnomeAD ^66,67^. We restricted our analysis to the 200 cancer missense mutations in positions that did not have a natural variant in any of the four human AGOs (Fig. 6b, Supplementary Data 6). We find that 87% of positions of these cancer-exclusive mutations belong to one of the signature positions, with most positions overlapping with the mAGO signature (60) and only a small number affecting universally conserved positions (16). Cancer-exclusive positions disproportionately affect the NA interface over population variants (Fig. 6c), albeit being localized more often in buried parts of the protein, as indicated by their mean solvent accessible surface (Supplementary Fig. 7b). Overall, we find that the cancer mutations impact on positions with a higher mean contact and sequence conservation (Fig. 6d), indicative of higher potential of possible deleterious consequences. Interestingly, a larger proportion of cancer-exclusive mutations cluster in the MID domain, when comparing the mutational distribution of variant and cancer mutations (Fig. 6e) and seem to be found more often in SSEs involved in NA binding (Supplementary Fig. 7c). In Figure 6f we highlight seven NA interface positions that are mutated in the cancer data (Fig. 6g) and the two in LESKRES syndrome (Fig. 6a).

In order to experimentally test our findings, we performed mutations on hAGO2 in human HEK293 cells. For this we deleted *AGO2* in the cells using a CRISPR/Cas9 approach. We then expressed selected mutant GFP-tagged AGO2 and measured protein stability, localization under stress and silencing effectivity. We selected a number of mutations based on signature profile, location in the protein and possible cancer evidence. Most of the mutants are located in the NA interface and the impact of mutating these amino acids are summarized in Fig. 6h. We find that almost all mutations in universal signature positions lead to failure in every functional assay, including very low protein stability in the cell (Supplementary Fig. 7d,e,f). Using co-transfection of a synthetic miR-21, we tested the ability of these mutant proteins to target a number of miR-21 target genes (Supplementary Fig. 7e). The silencing effectiveness was substantially reduced in all tested mutants when compared to WT hAGO2, except for the highly conserved eAgo signature position C.L14.1 which shows comparable levels of silencing. Interestingly, we also find two cancer mutations (C.L18.5 E673K, N.H7.2 D358G), where silencing is retained for selective genes. Indeed, when mutating the same positions to the standard alanine instead of the cancer mutation, silencing is abolished more uniformly across the miR-21 target genes (Fig. 6i), highlighting the specific effect of the cancer mutation. Further to this, hAGO2 immunoprecipitation was performed on hAGO2 mutant/ miR-21 co-transfections and we performed qRT-PCR on miR-21 target RNA isolated from the immunoprecipitates. This revealed an enrichment of a set of mRNAs from known cancer genes, most strongly in the N.H7.2 D358G mutant, including *PDCD4* and *PKD2* (Fig. 6j). This enrichment was less pronounced in the N.H7.2 D358A mutant. Finally, we tested the ability of each mutant to localize to stress granules following Sodium Arsenite exposure (Fig. 6k, Supplemental Fig. 7f, summarized Fig. 6h). While the majority of mutants tested demonstrated a severely abrogated capacity to localize to granules, N.H7.2 D358G and N.H7.2 D358A substantially retained WT-like stress localization. Our data reveals an accumulation of cancer mutations in the NA interface and provides novel insights into how these mutations can impact on NA interaction and oncogenic potential of Argonautes.

### The putative nucleic acid binding interface of Med13 has diverged drastically

Based on recent structural findings, the yeast CKM component Med13 was classified as a new Argonaute sub-family (Supplementary Fig. 7g) ^41^. Med13 resembles an Argonaute with a PIWI, MID, N, L1, L2, and PAZ domain architecture (Fig. 7a). However, Med13 also contains a 500 amino acids long intrinsically disordered region (IDR) within the L2 domain, which is absent in Argonautes. A short segment at the end of that Med13-specific L2 domain occupies a large part of the central channel that corresponds to the NA interface in Argonautes. The authors noted that a regulated release of this blocking L2-segment may restore Med13 NA-binding ability and that the captured structural conformation may represent an autoinhibited state. However, despite the similarity to the overall Argonaute fold, it is still unknown if Med13 displays any functional Argonaute-like features such as NA-binding or target matching.

**Fig. 7.**
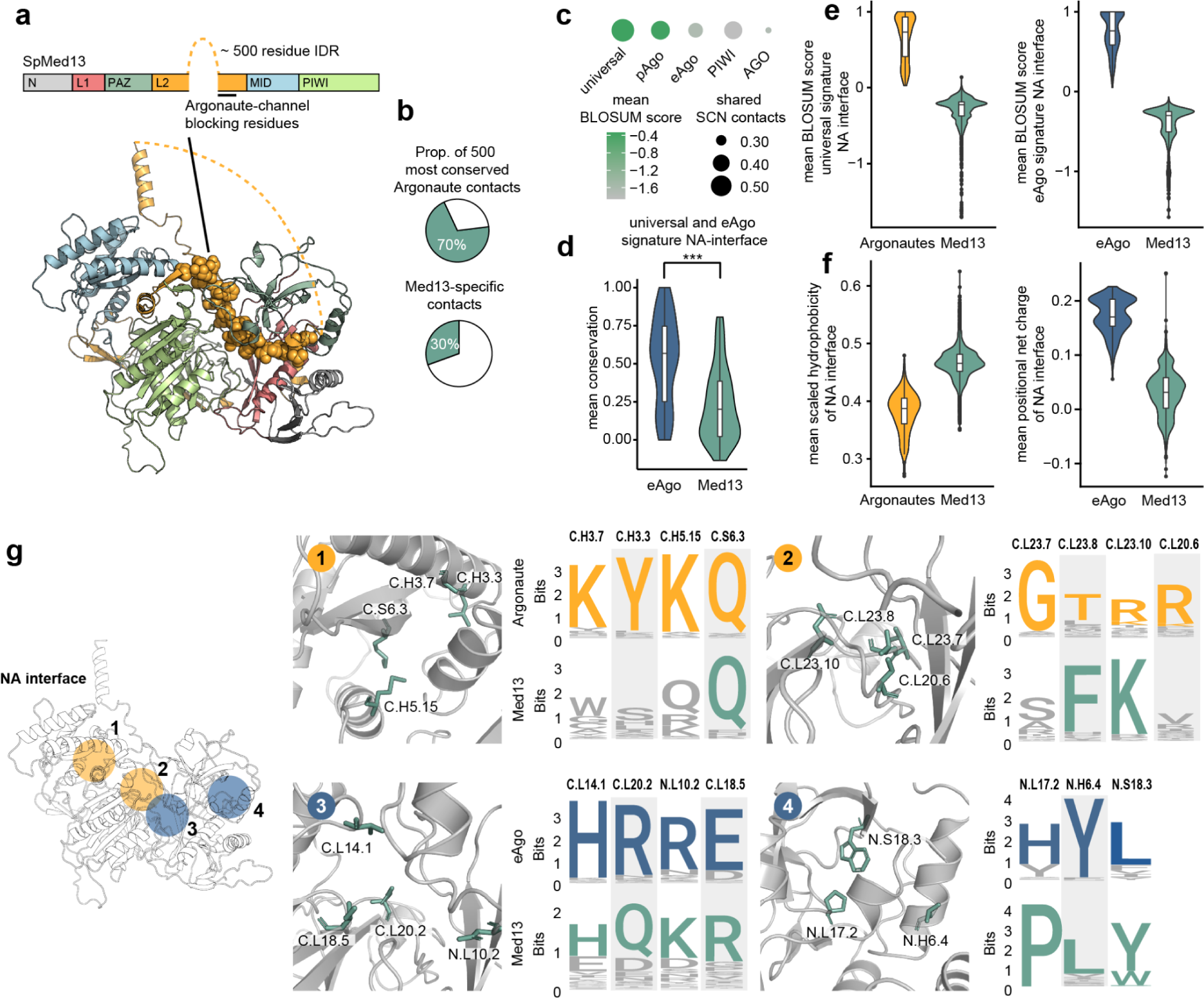
Comparison of the Med13 and Argonaute protein families. **a** Domain architecture and structure of the yeast Med13 protein (PDB: 7KPV). The Med13-specific L2 domain blocking the channel is highlighted as spheres in the structure. **b** Percentage shown for the 500 most common Argonaute contacts that are present in the yeast structure and the proportion of Med13-specific contacts. **c** Comparison of Med13 structure and sequence data with the universal, pAgo, eAgo, PIWI-clade and AGO-clade signatures and corresponding SCNs. The circle size represents the proportion of shared contacts with the respective SCN and the color represents the average normalized BLOSUM score against all Med13 protein sequences for the respective signatures. **d** Mean conservation of universal and eAgo NA interface positions in eAgo and Med13. **e** Normalized BLOSUM score of universal NA interface signatures against all Med13 and Argonaute protein sequences. eAgo NA interface signatures against all Med13 and eAgo protein sequences. **f** Violin plot of the mean scaled hydrophobicity and positional net charge calculated for all Med13 proteins and for eAgos and all Argonautes, respectively. NA interface is defined as shared pAgo and eAgo NA interface positions. **g** Examples of highly universal (orange) and eAgo (blue) signature NA interface residues and their corresponding positions in Med13 (most conserved residue highlighted green in logo representation).

To compare the molecular features of Argonautes and Med13, we constructed a multiple sequence alignment of Med13 across eukaryotes. We then cross referenced the positions in the alignment to the Argonaute topology (Supplementary Data 7). This enabled us to compare and analyze the contact network together with sequence conservation patterns between the Argonaute and the Med13 protein families. We find that the overall structural network of yeast Med13 contains 70% of the 500 most common Argonaute contacts, but that 30% of all Med13 contacts are not present in any other Argonautes from our set (Fig. 7b). When matched against the respective signature residues and SCNs, Med13 shares most contacts and sequence similarity with universal and pAgo signatures and SCNs (Fig. 7c).

The eAgo NA interface positions show a significantly lower conservation in Med13, hinting at a more variable channel (Fig. 7d). Indeed, we find that the universal and eAgo signature NA interface positions have diverged drastically in Med13, with low overall BLOSUM sequence scores for both (Fig. 7e). In consequence, this region is significantly more hydrophobic in Med13 and is less positively charged than the eAgo-channel (Fig. 7f). In Figure 7g we highlight selected universal and eAgo signature positions and their equivalent positions in Med13 across the NA interface, revealing differences in the universally conserved 5’MID and seed binding region, and the eAgo-specific NA-binding positions near the catalytic region and the PAZ domain. Our data indicates that while Med13 retains many of the universally conserved features of an Argonaute protein, it is unlikely to retain a capacity to bind nucleic acids through an Argonaute-like interface.

## Discussion

In this study, we present a comprehensive analysis of the common and diverging molecular features of Argonautes across extensive evolutionary timescales. Integrating structural information, phylogenetic data and sequence conservation allowed us to discern universal from clade-specific features in Argonaute evolution (Fig. 8a,b). We determined the signature positions for each subgroup and analyzed their local structural environment via contact networks. Using experimental structures we identified the universal and clade-specific NA interface signature positions. We validated our approach by performing mutational assays on selected signature positions in the hAGO2 NA interface, which resulted in impaired or reduced silencing efficiency, localization under stress and overall protein stability.

**Fig. 8.**
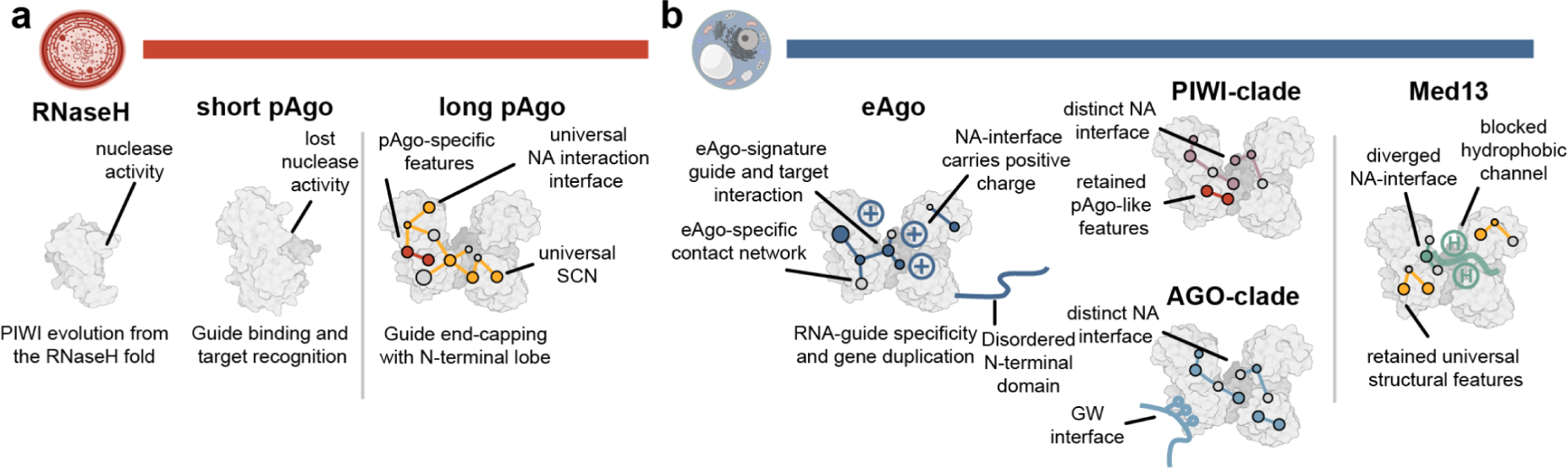
Summary for Argonaute structural evolution. **a** The Argonaute PIWI domain evolved in early prokaryotes from the RNaseH fold. Long pAgos adopt the typical bilobed structure, which is supported by the universal SCN and contains some pAgo-specific features. **b** Newly emerging structural features in eAgos range from a positively charged NA interface to a disordered N-terminal terminus. The PIWI and AGO-clade display distinct NA interface features, especially in the guide seed region. The PIWI-clade retains pAgo-like features and AGOs evolved multiple GW-binding pockets. Med13 adopts an Argonaute-like fold but displays a highly divergent NA binding channel.

Since universally conserved residues that form the basis for the seed NA interface are shared between RNA and DNA-guided Argonautes, it is unlikely that they are functionally relevant for guide-selection. Indeed, we find that many newly emerging eAgo signature positions contact the second half of the guide, which is positively charged when compared to pAgos. *In vitro* experiments have shown hAGO2 ability to bind DNA guides for targeting albeit with lower efficiency and the additional C2’ OH group were proposed to play a role in guide tolerance ^50,68^. DNA/RNA hybrid-guide experiments have shown that changing the 5’ half of guides from RNA to DNA led to no substantial loss of gene-silencing activity, while the 3’ half was more sensitive to DNA-substitution ^51^. Thus, we speculate that eAgo RNA-guide preference is mediated by eAgo-signature residues in the N-Lobe that contact the RNA ribose backbone C2’ OH directly. We identify candidate positions that satisfy these conditions, but further biochemical experiments are needed to study their impact on NA binding. Importantly, we cannot exclude that more complex conformational differences between RNA and DNA (e.g. ribose sugar puckering) play a decisive role in the guide selection and duplex structure.

We find that MfAgo has sequence features that closely resemble an Argonaute of eukaryotic origin and has a positively charged NA interface. However, the MfAgo structural model displays only few of the contacts typical of eAgos. This, together with its low similarity to any other pAgo, may suggest that it represents a close relative to eAgos that has diverged during evolution. Previous studies have supported an archaeal origin of eAgos ^46,69,70^. However, the similarities may have also occurred due to convergent evolutions as, for instance, the thermophile may rely on high interface charge in MfAgo to stabilize the NA interaction.

Since the AGO- and PIWI-clades are broadly represented across major groups in eukaryotes, it was previously proposed that a duplication event likely predated the eukaryotic common ancestor ^46^. Here, we identify the distinct structural and sequence signatures for both clades. Beyond the AGO-specific central gate positions, many of the diverging signature positions between the PIWI and AGO-clade NA interface are located in the seed-binding region, suggesting distinct properties of this region. Recent work has revealed that PIWIs do not pre-organize the full piRNA seed region and that PIWI seed complementarity is dispensable for target cleavage ^34,37^. Our analysis identifies conserved changes in the PIWI-guide seed interface that differs from the AGO-clade and may impact on guide affinity and target matching. Especially the conserved changes to helix N.H7 are plausibly consequential in this regard, as mutations were shown to impact on target selection by increasing the binding to targets with imperfect seed complementarity ^71^.

Comparing and scoring PIWIs and AGOs against all pAgo proteins included in our study revealed that the PIWI-clade-specific features show higher agreement with most pAgos. In addition, we identify proteins in archaeal and bacterial pAgo branches that share distinct sequential and structural signatures with the PIWI-clade. Our results are supported by previous studies that found functional similarities between pAgos and the PIWI-clade, for example the positioning of the glutamate finger in the active site and the metal-mediated 5′-guide interactions in the MID pocket. Interestingly, the role of PIWI-clade targeting against evolving mobile genetic elements resembles in some aspects the pAgo host-defense function: Both require enough flexibility to accommodate rapidly evolving target sequences and enough stringency to avoid off-target effects. In principle, these similarities thus could have emerged via processes of convergent evolution. However, we also find similarities in regions of low structural significance that are more likely to stem from homology.

Our analysis supports a model in which Argonaute GW-binding sites accumulated during AGO-clade evolution. In hAGO2 the three identified TRP-binding pockets were shown to not be fully redundant with each other, since individual GW-motifs showed preference for specific pockets ^62^. It is thus plausible that the emergence of additional pockets within the AGO-clade allowed for a more tunable Argonaute-GW-protein interaction in a cellular environment where multiple GW-motif-carrying proteins may compete for binding. Argonaute interaction with GW/WG motif-carrying proteins have been reported in animals, plants and yeast. Thus, they were suggested to be a conserved motif in otherwise unrelated Argonaute-interacting proteins ^57,59,72^. Interestingly, the motif was also found to be exploited for RNA silencing suppression by the *Sweet potato mild mottle virus* ^73^. Due to the relative simplicity of the motif, convergent evolution of interactions facilitated through disordered segments likely drove functional specialization.

Mutational data of hAGOs support the importance of many of the signature positions identified in this study. We find that cancer mutations accumulate in the NA interface and may impact binding directly. Many studies have focused on the dysregulation of small RNAs in tumorigenesis and an increasing number of miRNAs have been identified as potential biomarkers for diagnostic and prognostic purposes ^74^. Depending on the biological context, miRNAs may function as oncogenes or tumor suppressors. In many cancers, Argonaute proteins are overexpressed, highlighting their oncogenic potential ^65,75,76^. Mutations in the NA interface may modulate or shift the sets of miRNAs that are bound or change selection criteria for putative targets. Indeed, in our mutational assay cancer position N.H7.2 D358G impacts hAGO2 silencing selectively and alters the binding preference for a number of genes related to cancer. Given the crucial role of hAGO proteins in the maturation of small RNAs, mutations in the NA interface could also affect their availability and homeostasis impacting on transcriptional control in cancer cells. More research on the effects of these mutations will be needed to identify their biochemical and biological consequences.

Finally, we applied our analysis to the CKM component protein Med13 to determine if the core characteristics are preserved in the structurally related protein. Due to the broad similarities to the typical Argonaute fold it was proposed that Med13 may have NA binding capability ^41^. We find that despite retaining many of the structural Argonaute properties, most putative NA interface residues have diverged substantially from the universal Argonaute signature. As a result, the Med13 channel is much more hydrophobic than that of the average Argonaute and the overall low conservation of the Argonaute channel may hint at co-evolutionary processes with the blocking L2 domain. Typically, NA backbone binding is conferred by residues that are well conserved across protein families because the NA epitope is stable across different species. This is in contrast to co-evolving protein-protein interfaces ^44,77^. Based on our results, we propose that the differences in the putative NA interface positions preclude an Argonaute-like binding mechanism. However, this does not exclude Med13 NA-binding through a different and potentially Med13-specific binding interface.

Our study redraws the evolution of Argonautes by separating universal from clade-specific features and integrating them with biological function. The signature positions and their contact networks determined in this study can provide starting points for new experiments to better understand Argonaute function and dysfunction in disease and provide a framework to identify Argonautes of biological interest. Our work can serve as a template for how to study structure in the age of advanced structure prediction tools by utilizing and integrating experimental structures and structural models to uncover patterns in the structural evolution of protein families.

## Methods

### Structure and AF2-guided Argonaute alignment

We used an iterative approach to build a structure-based alignment: First, all available Argonaute structures were collected from the PDB ^78^ (85 structures, Supplementary Data 1) and one structure per organism was selected for an initial structure-based sequence alignment with MUSTANG ^79^. The selection of structures was based on equivalent binding state (guide binding) and overall quality (resolution, completeness, mutations). The following structures were selected for the alignment: 4F1N, 4F3T, 6OON, 5VM9, 4KRE, 5GUH, 6KR6, 7KX7, 7SWF, 3DLB, 6QZK, 5AWH, 5I4A, 1YVU, 7R8K, 5G5T, 1Z26). Residues missing from the structure and mutated residues were included without changing the underlying alignment using MAFFT ^80^. With this initial alignment, the profile-based JACKHMMER ^81^ was used to search the entire UniprotKB database ^82^ with a significance E-value cutoff of 0.01 (Sequence), 0.03 (Hit) and 5 iterations. Next, the resulting matches were first filtered by length (70% of query length) and sequence similarity to reduce the initial number of hits. To ensure broad sequential diversity and a sufficient coverage of the phylogenetic tree and Argonaute clades in eAgos, we selected Argonaute-representatives according to the UniProt taxonomy database annotations. Representation of pAgos was guided by taxonomy information derived from the Genome Taxonomy Database (GTDB) ^83^ database, using AnnoTree ^84^ for visualization. After the selection of the sequences, the structures of the segments that matched to the query alignment were predicted with AlphaFold2 ^42^ using ColabFold ^85^ or, if available, downloaded from the AlphaFold database ^86^. The structural models were inspected visually to confirm fold and homology. To increase the accuracy of the structure-guided topology the 10 best-scoring (mean plDDT) structural models from Eukaryota, Archaea and Bacteria were structurally aligned together with the above selected structures from the PDB to create a new multiple sequence alignment, which then was used to search the UniprotKB database a second time using JACKHMMER in the manner described above. Newly identified matches that qualified the filter criteria laid out above were added to the selection. In a final step the selection of sequences were added to the query structure-based sequence alignment using the MAFFT *–add* method ^80^ (FFT-NS-2, BLOSUM62 scoring matrix) to preserve the initial structural alignment as much as possible. Using the final reference alignment the full UniprotKB was searched a third JACKHMMER search yielding 17,461 putative Argonaute-like sequence matches.

### Argonaute topology

Based on the consensus secondary structure of the human Argonautes we devised an Argonaute topology assigning each position in the alignment. The topology was expanded to the alignment to assign a unique structure-based identifier for each position. The topology identifier combines information on the lobe (C or N), the structure element (H for helix, S for strand and L for loop). These structural elements are numbered from their appearance within the respective lobe (e.g. C.H1 represents the first helix of the C-lobe). Finally, the last digit identifies the position within that secondary structure element (e.g. C.H1.2 represents the second position in the aforementioned helix). Another digit is added if there is no corresponding hAGO position in the alignment (e.g. C.H1.2.1). A labeled map of the Argonaute topology can be found in Supplementary Fig. 1f, which was created using Pro-origami ^87^. Supplementary Data 2 lists all amino acid sequence positions for all topology positions of the proteins in the reference alignment.

### Sequence conservation and signature positions

For any given group we extracted the sequences from the alignment based on phylogenetic considerations or sub branches/clades within reconstructed Argonaute tree and calculated the normalized conservation score (NCS) for each position in the alignment, the mean conservation score (mNCS) and the standard deviation (sdNCS) of all positions. We considered a position *p* as candidate of the respective signature if NCS*_p_* > (mNCS + 2 * sdNCS). We then conducted a pairwise comparison of the respective signature candidates to all higher-order signatures that the group may also belong to (e.g. the metazoan AGO-clade will have signatures that emerged in the AGO-clade, eAgos and universal signatures) and excluded all position that have been conserved as the same residue. However, if a given signature candidate *p* is divergent from any of the higher-order signatures, those positions were scored pairwise against the higher-order signatures using the normalized BLOSUM matrix (pBS). We define a sufficiently changed residue if pBS ≤ 0. If a position met this criteria it was included in the respective signature, otherwise the position was excluded. In order to score a set of signature positions with individual proteins we calculated the mean pairwise score pBS of each signature position against the equivalent position of a given sequence yielding one average score per analyzed sequence/protein.

Because the number of sequences and average sequence conservation in the reference alignment differs between the Archaea, Bacteria and Eukaryota kingdoms, we chose an approach that was recently applied on the TATA-box binding protein (TBP) ^44^ to identify the universally conserved residues in Argonautes. Briefly, we reconstructed the ancestral sequences for bacterial, archaeal and eukaryotic Argonautes using FASTML ^88^ using the sequences from the reference alignment. The resulting ancestral sequences were referenced to the topology via the reference alignment by deploying the MAFFT *–add* method like above. The added sequences did not change the alignment size. Thus, each position was scored among the ancestral sequences by calculating the normalized BLOSUM (BLOSUM62) score for each position as described in detail in Ravarani et.al ^44^. This was combined with an additional minimum normalized conservation threshold of NCS*_p_* > (mNCS + sdNCS) to validate the overall conservation of the positions across the whole data set. Normalized similarity conservation scores were calculated using the *conserv* function from the Bio3D R package with a normalized BLOSUM62 substitution matrix ^89^.

Sequence net charge was determined using the Henderson-Hasselbalch Equation (pH7) and averaged by position (positional charge). Mean scaled hydropathy was calculated using a normalized Kyte and Doolittle scale.

### Intra- and intermolecular interaction data

Residue to residue salt bridges, pi-cation, pi and t-stacking, hydrogen bonding and van der Waals contacts were determined using the GetContacts application (https://github.com/getcontacts/getcontacts). These contact networks were calculated for all available structures in the PDB and all predicted structures from this study. AF2 contacts were only considered when both contact partners had an AF2 pLDDT >= 80. Integrating the structural topology naming scheme, sequence conservation and structural information was realized with custom R scripts.

Intermolecular contacts between the protein and the nucleic acid interface were determined in an equivalent manner. To enable comparison of NA-positions in a structurally meaningful way, we referenced all positions to a common naming scheme (illustrated in Supplementary Fig. 3e). Some Argonautes have different guide lengths, but the 3’ PAZ domain anchoring remains comparable. Hence, in order to compare those contacts in a position-specific manner, the last 3 positions of any guide are counted from the 3’ end (3T, 3T-1, and 3T-2 in Supplementary Data 5). We then manually referenced all target strand positions in the available structures to its complementary guide position (e.g. the target position that matches with the 5th guide position is named -5). We excluded mismatched and bulged positions from our analysis, but they are included in the provided data and are marked with “B” together with the number they would ordinarily match with (e.g. B-5 mismatches with guide position 5). For both the universal and eAGO signature NA interface we only included positions that contacted the same guide/target position in at least one pAgo/eAgo or PIWI/AGO representative, respectively.

### Contact conservation and SCNs

Each Argonaute structure/model yielded a network of pairwise contacts between topology positions. This enabled us to analyze each node (topology position) and its contacts (edges) across groups of proteins. Contact conservation (CC) reflects the presence of a contact within a defined group (e.g. if a contact is present in 50% of all eukaryotic structures/models, the contact conservation is 0.5 in eAgos for this contact). The mean contact conservation of any given position or region is defined as the mean contact conservation of all contacts that position/region is involved in.

To determine the SCN for a subset of proteins, we calculated the contact conservation for each contact in the alignment and determined the mean contact conservation (mCC) and the standard deviation (sdCC). We then considered a contact *c* as part of the respective SCN if CC*_c_* > (mCC + sdCC) and at least one of the two nodes of the respective contact is part of the respective signature set. For the calculation we only considered one structural representative per protein, if more than one structure was available for one protein. Because pAgos are more structurally variable than eAgos, we only considered a contact part of the universal SCN if CC*_c_* > (mCC + sdCC) was true for both the eAgo and pAgo subgroups.

### Visualization and figures

PyMol (The PyMOL Molecular Graphics System, Schrödinger, LLC, Version 2.5.2) was used to manually evaluate structures and create the structure images for the figures. Contact networks were visualized with the ggraph R package. The sequence logo plots were created with the R package ggseqlogo ^90^. The phylogenetic tree was plotted and any heatmap annotation of the tree was created using the R package ggtree ^91^.

### Phylogenetic tree construction

Based on the structure-guided alignment an initial maximum-likelihood tree was constructed with FastTree, which was subsequently used for further maximum-likelihood optimization with the MEGAX software ^92^ (WAG (g+I)) yielding the final tree.

### Statistical testing

All significant differences reported here were tested with the Wilcoxon rank-sum test with significances noted as following * p < 0.05, ** p < 0.01, *** p < 0.001.

### Disorder propensity

IUPred2A was used to estimate disorder propensities of the N-termini of Argonaute proteins^93^. The N-terminus was defined as every position preceding the JACKHMMER match that was used for the reference alignment. The prediction was done for each protein sequence individually using the R package idpr ^94^ and the mean and standard deviation was subsequently calculated and plotted with R. A disordered propensity value of >0.5 for a given position can be considered disordered.

### Mutational data

Natural variation and cancer mutation data for all four human AGO-clade Argonautes were obtained from the gnomAD database and the CBioPortal, respectively. LESKES syndrome hAGO2 mutations were taken from Lessel et. al ^64^. All missense mutations were translated into the Argonaute topology positions via the reference alignment. Cancer-exclusive mutations were defined as affecting topology positions that did not have any natural variation mutation in any of the human AGOs. The resulting positions were annotated by SSE, signature (vAGO, mAGO, AGO-clade, eAgo and universal) and nucleic acid interface positions. The distribution of cancer and variation mutations for each domain was determined by calculating the mutation distribution normalized by domain size and number of available missense mutations in each dataset.

### Med13 alignment and analysis

We searched the Representative Proteomes (RPs) 55 UniprotKB database with the yeast Med13 sequence (UNIPROT: P38931) using phmmer and the same settings, repeats and query-length cutoff like in the Argonaute sequence search above. The search yielded 1939 sequence matches exclusively from eukaryotes, from which we calculated a multiple sequence alignment using MAFFT. We then structurally aligned Med13 (PDB:7KPV, chain:D) using Mustang to reference the structure to our Argonaute topology. We were thus able to extract the structurally equivalent positions from the MSA and analyze and compare selected positions between Med13 and Argonautes (Data table 7). The intermolecular contacts of the 7KPV:D structure were calculated in the manner described above and referenced via the Argonaute topology.

### Cell culture

Hek293 cells were cultured in Dulbecco’s Modified Eagle Medium (DMEM, Gibco) containing 10% Fetal Bovine Serum (A5256701, Gibco), 1% Pen/Strep and 1% NEAA. Cells were passaged every 2-3 days, media was removed and cells were washed with 1X PBS (Gibco). Once PBS was removed, cells were detached through 3 minutes incubations with 0.05% Trypsin-EDTA, neutralised with 2X volume cell culture media and centrifuged for 3 mins at 1200RPM. Pellets were resuspended in complete media and plated at appropriate confluency on tissue culture treated flasks or plates (Gibco).

### CrispR deletion of *AGO2*

In order to generate *AGO2*^-/-^ Hek293 clones, cells were passaged and plated on 24-well plates (Corning) in antibiotic free conditions at 30% confluency. Per well, 1µM *AGO2* guide RNA (Hs.Cas9.AGO2.1.AB,Integrated DNA Technologies) was incubated with 1µg TrueCut Cas9 v2 (Thermo Fisher) in 100µl OptiMEM (Gibco) for 5 minutes at 37°C. Upon formation, complexes were mixed with 1µl Lipofectamine 2000 (Thermo Fisher), which was prepared in 100µl OptiMEM, and incubated for 20 minutes at 37°C. Liposome complexes were added to cells and incubated O/N. The following morning, cells were refreshed with media containing antibiotics and cells were incubated for a further 24hrs. Individual colonies were hand picked and grown in individual wells, prior to expansion and genotyping. Confirmed mutant clones were expanded and used for further analysis.

### *AGO2* mutant plasmid generation

EGFP-hAGO2 was a gift from Phil Sharp (Plasmid # 21981, Addgene). Using Phusion Site-Directed Mutagenesis kit (F542, Thermo Fisher) and individual 5’ phosphorylated primer sets, 16 mutant *AGO2* plasmids were generated. 10ng of wt plasmid was used as template in each reaction, and PCR was conducted according to manufacturer’s protocol, with annealing temperature set as shown in Primer Table. Upon completion, 5µl of product was run on 0.75% agarose gel to confirm successful amplification. Following this, DpnI digestion was performed by adding 1µl of FastDigest *DpnI* enzyme to the PCR reaction and incubated at 37°C for 15 mins. Ligation was performed according to manufacturer’s protocol using 5µl of *DpnI* digested PCR product and upon completion, samples were stored at -20°C prior to transformation. Product was transformed into *DH10-B* Competent Cells (F542, Thermo Fisher) according to manufacturer’s protocol and grown up O/N on Agar plates with Kanamycin selection. The following day, individual colonies were picked and further grown in 20ml LB Broth (10855001, Gibco) containing Kanamycin. Plasmid DNA was purified using a QIAprep Spin Plasmid MiniPrep kit (27104, Qiagen), as per manufacturer’s protocol.

### *AGO2* plasmid transfection

*AGO2^-/-^* Hek293 cells were plated at 70% confluency on 24-well plates in antibiotic free conditions, as described above. Per well, 1µg of each mutant plasmid was mixed with 1µl of Lipofectamine 2000, in 200µl OptiMEM, and incubated for 20 minutes at room temperature. Upon completion, mixtures were added to appropriate cells and incubated overnight. The following morning, media was refreshed and cells were further incubated. For miR based assays, cells were incubated for 24 hours prior to further treatment. For cycloheximide assay and live cell imaging, cells were incubated for 48 hours prior to further treatment.

### *miR-21* transfection

24-hour transfected *AGO2^-/-^* Hek293 cells, as described above, were prepared for each mutant clone. 5pmol mirVana human miR-21 (Ambion) was mixed with 1µl of Lipofectamine 2000, in 200µl OptiMEM, and incubated for 20 minutes at 37°C. Upon completion, mixtures were added to cells and incubated overnight. The following morning, media was refreshed and cells were further incubated for 48 hours total, prior to harvesting.

### Live cell imaging and stress granule assay

48-hour plasmid transfected *AGO2^-/-^* Hek293 cells, as described above, were prepared for each mutant clone on 12-well plates (Corning). Prior to imaging, media was refreshed, containing 1µg/ml Hoechst 33343 (Thermo Fisher), and incubated for 20 minutes at 37°C and 5% CO_2_, after which cells were returned to standard media. Cells were placed in the live cell imaging chamber attachment in ImageXpress Pico (Molecular Devices) with the environment maintained at 37°C, 80% humidity and 5% CO_2_. Cellular GFP (500 msec) and Hoechst (30 msec) activity was imaged for each clone. In order to induce AGO2 localisation to stress granule as has been previously demonstrated ^95,96^, following imaging, cells were exposed to media containing 100µM Sodium Arsenite (Merck), and incubated for 40 minutes 37°C and 5% CO_2_. Cells were re-imaged in the ImageXpress Pico following incubation, as above.

### Cycloheximide exposure

48-hour plasmid transfected *AGO2^-/-^* Hek293 cells, as described above, were prepared for selected mutant clones on 12-well plates (Corning). Media was refreshed, containing 100µg/ml cycloheximide, and cells were further incubated for 24 hours.

### AGO2 immunoprecipitation (IP)

48-hour plasmid transfected *AGO2^-/-^* Hek293 cells, as described above, were prepared for selected mutant clones on 12-well plates (Corning). Cells were washed with ice cold 1XPBS and lysed with ice cold Pierce IP Lysis Buffer (Thermo Fisher) containing cOmplete protease inhibitors (Merck), according to manufacturer’s protocol. Lysate was transferred to a 1.5ml tube and centrifuged at 13000RPM for 10 mins at 4°C. Supernatant was transferred to a fresh tube and used for immediate processing. 25µL Pierce Protein A Magnetic Beads per sample was transferred to a 1.5ml Eppendorf and diluted with 150µL Wash Buffer (Tris-buffered saline containing 0.05% Tween-20). Immunoprecipitation was performed following manufacturers protocol, with minor modifications. A magnetic separation column (NEB) was used to isolate beads and wash solution was removed. Protein lysate was added to mixed with beads and 2µL Argonaute2 antibody (C34C6, Cell Signaling), and rotated at 4°C for 2hrs. Following antibody binding, magnetic separation was performed and remaining solution was removed. Samples were washed three time in ice-cold Wash buffer with 5 mins of rotation at 4°C, prior to magnetic separation and removal of buffer. Upon completion, bound samples were either processed for protein or RNA, as described below.

### RNA extraction, Reverse Transcription and RT-QPCR

Samples from either the IP Assay or the miR transfection assay were processed for determining gene expression levels. RNeasy Mini kit (Qiagen) was used as per manufacturer’s protocol without modification. Samples were eluted in 15µL RNase free H_2_0 and concentrations were determined. 500ng of RNA was used for reverse transcription. SuperScript III (Thermo Fisher) reverse transcriptase was used as per manufacturer’s protocol, with oligo(dT) primers used for amplification. Samples were incubated at 65°C for 5 mins, followed by 50°C for 50 mins and finally 75°C for 15 mins to terminate the reaction. Newly generated cDNA was diluted with clean H_2_0. For RT-QPCR analysis, each reaction was performed in triplicate wells and each plate run in duplicate. Power SYBR Green Mastermix (Thermo Fisher) was used according to manufacturer’s protocol in 10 µL volumes, with 2 µL of diluted cDNA used per well. Plates were run on a Quant Studio 6 Flex (Applied Biosystems) and data was analysed on Quant Studio Flex software.

### Western Blotting

Samples from either the Cycloheximide Assay or the IP were processed for determining AGO2 levels. For cycloheximide assay, 1X RIPA Lysis buffer (Thermo Fisher) containing cOmplete protease inhibitors was added directly to cell pellets. Following complete lysis, samples were centrifuged at 13,000RPM for 10 mins at 4°C to pellet cell debris. Supernatant was removed to a clean tube and an equal volume of 2X Laemmli Buffer (Bio-Rad) was added, prior to incubating samples at 95°C for 5 mins. For the IP samples, 2X Laemmli Buffer was applied directly to the washed beads and incubate for 5 mins at RT, prior to incubating at 95°C for 5 mins. Samples were loaded onto Mini Protein TGX 4-15% gels (Bio-Rad) with PageRuler Prestained Ladder (Thermo Fisher) used with each gel. Gels were run for 90mins at 100V, or until sufficient separation had been achieved. Protein was transferred to nitrocellulose membranes in Tris/Glycine-Methanol buffer by running at 100V for 45 mins. Membranes were briefly washed in 1X TBS-T, prior to blocking in blocking buffer (5% skimmed milk in 1XTBS-T) for 1hr at RT. Primary antibody incubation was performed with Argonaute2 antibody (C34C6, Cell Signaling, 1:1000) in blocking buffer at 4°C with rotation O/N. The following morning, membranes were washed 3X in 1X TBS-T for 5 mins with rotation, prior to secondary antibody incubation (Alexa Fluor Plus 555-conjugated Goat anti-Rabbit, A32732, Thermo Fisher, 1:3000) in blocking buffer for 3hr at RT with rotation. Upon completion, membranes were washed 3X in 1X TBS-T for 5 mins with rotation, prior to imaging with ChemiDocMP Imaging System (Bio-Rad). For IP blots, exposure was 5 secs, while for cycloheximide blots, exposure was 12 secs.

## Primer Table

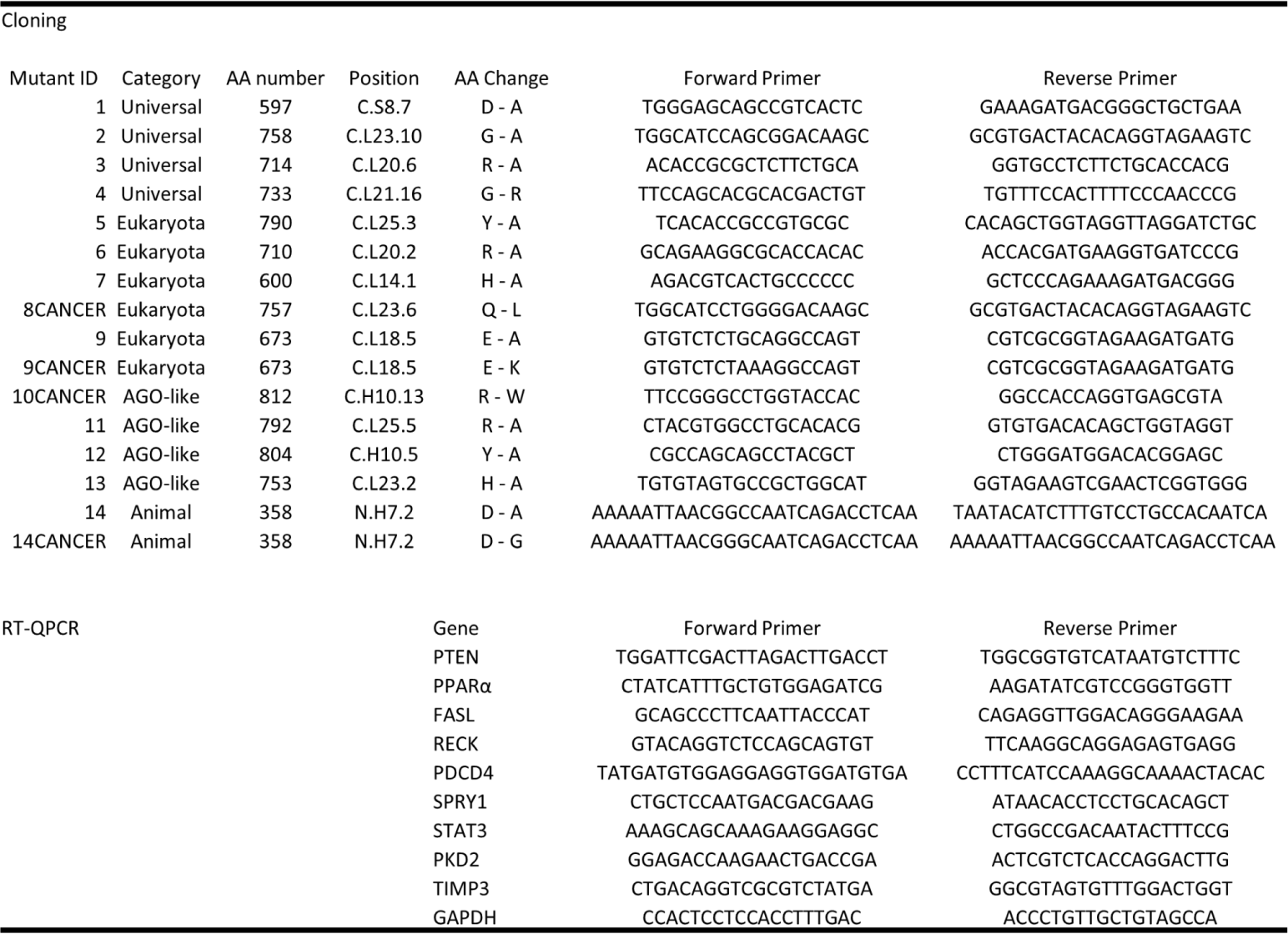

## Data availability

The data that support this study is provided as supplementary data files.

## Acknowledgements

We thank Camilla Ventura Santos (Max-Planck-Institute for Biophysics, Frankfurt), Daan Swarts (Wageningen University & Research), Emma Hands (MRC Laboratory for Molecular Biology, Cambridge) and Lajos Kalmar (MRC Toxicology Unit, Cambridge) for discussions. Logos for pAgos and eAgos in Figures 8a,b were created with BioRender.com.

## Author contributions

A.W. and M.V.D.P. designed the research. A.W. performed the research and analyzed data. M.V.D.P. performed and analyzed the mutational experiments. A.W. and M.V.D.P. wrote the manuscript.

---

**Supplementary Fig. 1.**
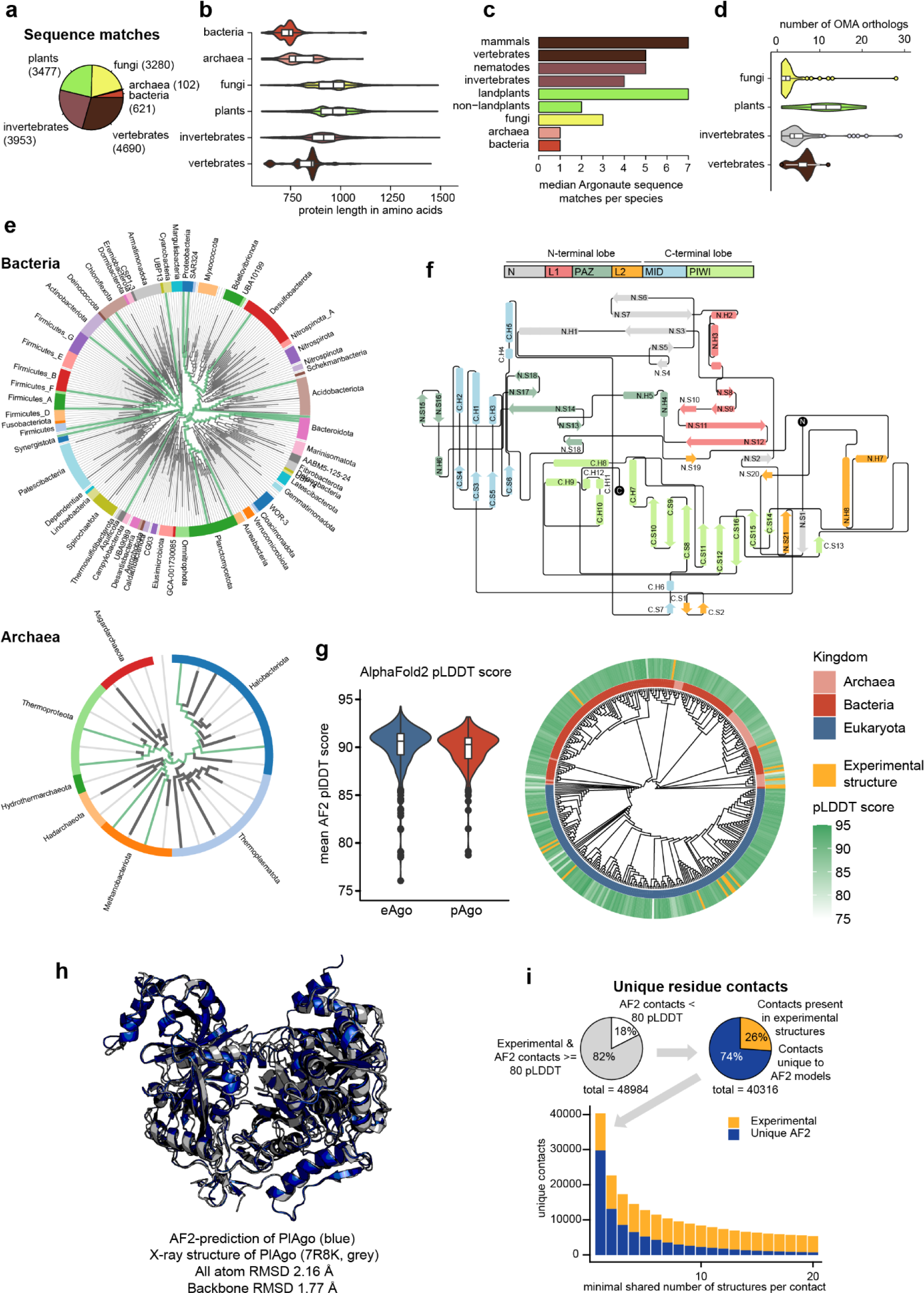
**a** Pie chart of phylogenetic origin of Argonaute sequences found in the UniprotKB database. **b** Violin plots of length of putative Argonaute sequences by phylogenetic origin. We set a cutoff at a length of 1500 amino acids for visualization purposes. **c** Median number of Argonautes per species found in this study. **d** Number of Argonaute proteins sourced from the OMA database across fungi, plants, invertebrates and vertebrates. **e** Bacterial and archaeal phylogenetic class annotation derived from the Genome Taxonomy Database (GTDB) database and visualized with AnnoTree. Taxonomies with pAgo representatives included in the reference alignment are highlighted in green. **f** Cartoon representation of the Argonaute topology with labeled helices and beta-strands. **g** Boxplot of mean full protein pLDDT scores from AF2 and projection on Argonaute dendrogram with experimental structures highlighted in yellow. **h** Structural overlay of AF2-predicted structural model of *Pseudooceanicola lipolyticus* Argonaute (PlAgo) and the published structure (PDB: 7R8K). **i** Full residue contact network from all available structures in this study. Contacts were only considered that were either from experimental structures or both contact partners had an AF2 pLDDT >= 80. Of the full set of contacts used in the study 26% can be found in at least one experimental structure. The proportion of contacts present in at least one experimental structure increases when filtering by the minimal shared number of contacts.

**Supplementary Fig. 2.**
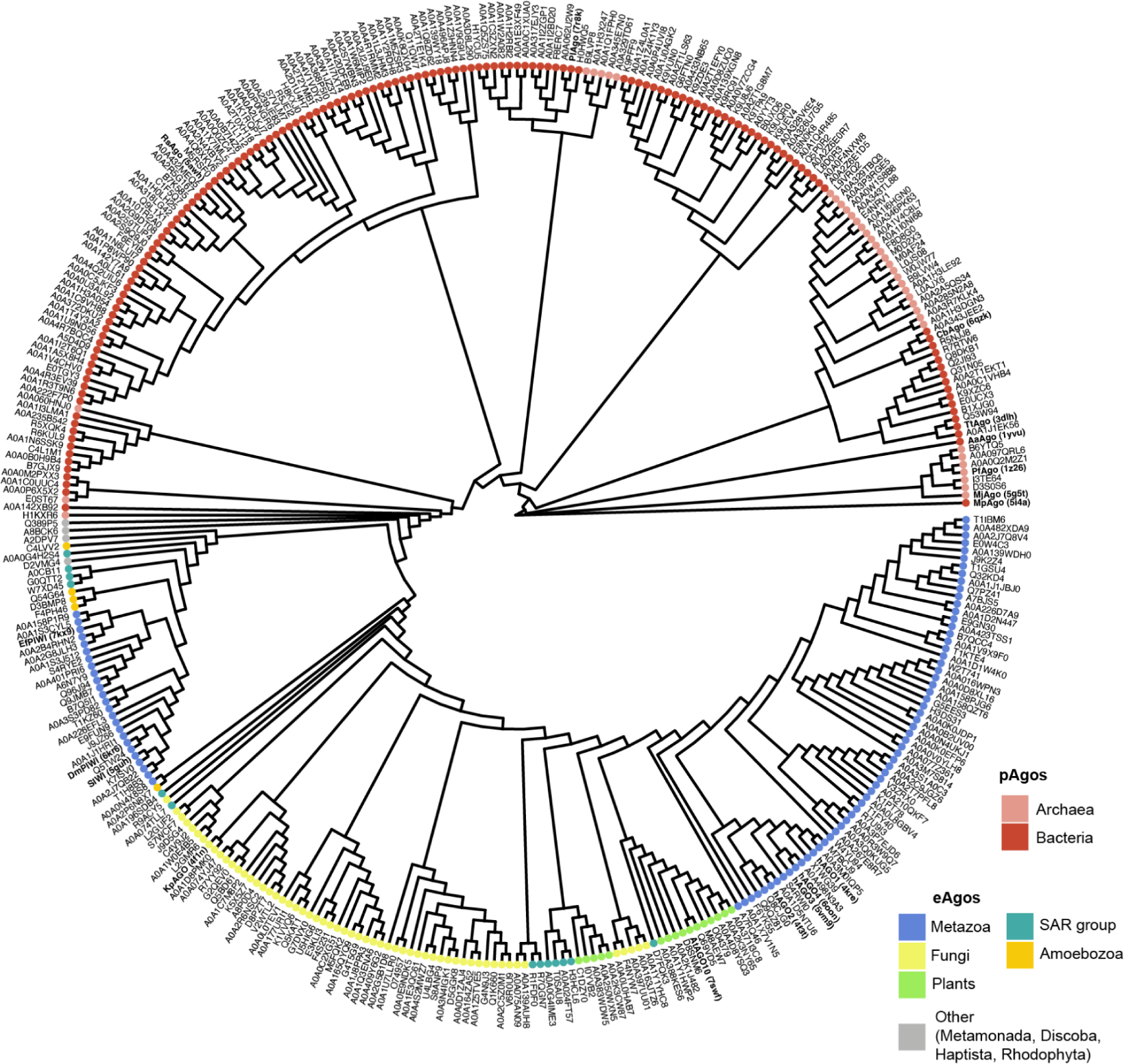
Detailed phylogenetic tree of all protein sequences used in this study. The nodes of the tree are color-coded by phylogenetic group and annotated with the UNIPROT codes. Proteins with available experimental structures are annotated in bold by name and PDB code of the published structure that was used for the structural alignment.

**Supplementary Fig. 3.**
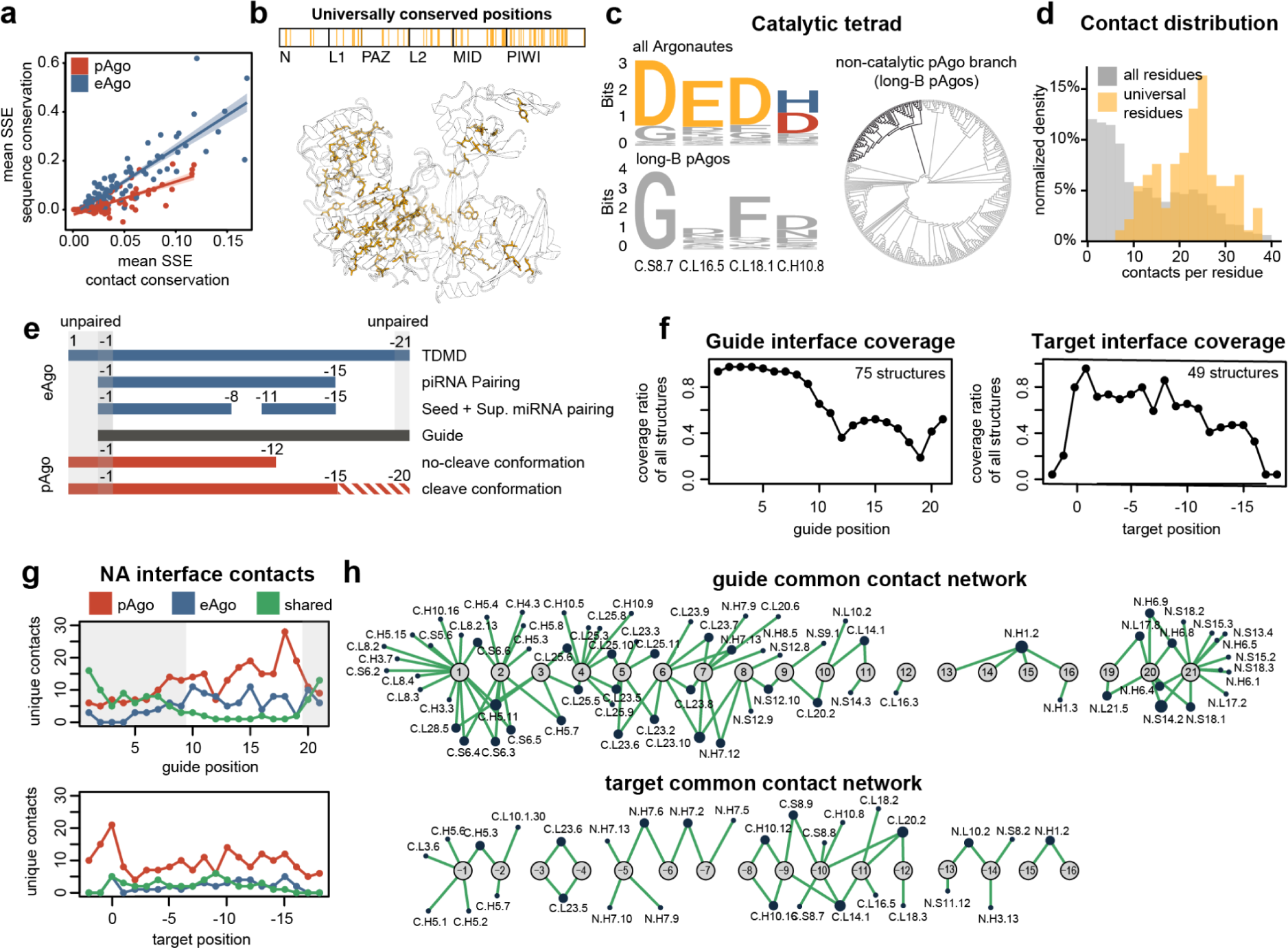
**a** Correlation plot of the mean contact and sequence conservation by SSE across all eAgos and pAgos. **b** Strip plot and cartoon structure representation of all universal signature positions. **c** Sequence logo plot of the catalytic tetrad for all Argonaute sequences and non-catalytic pAgos (long-B pAgos). Universal (yellow), pAgo (red) and eAgo (blue) -specific signature residues are highlighted. The long-B Ago branch is highlighted in the phylogenetic tree. **d** Histogram of contacts per position for all (grey) and universal signature residues (yellow) across all structures and models. **e** Guide and target numbering scheme employed in this study: Guide numbering 1-21 (for referencing of longer guides see Methods). The target was referenced to its complementary guide positions (e.g. target position -5 is complementary to guide position 5, positive target positions are unmatched 3’ target ends). **f** Coverage of each guide and target position in all available structures that include either guide or both guide and target chains. **g** Total number of unique shared (green) and eAgo (blue) or pAgo (red) -specific contacts resolved by guide (top) and target (bottom) position. Guide contacts are predominantly shared in the seed and 3’ end binding region (highlighted in grey). **h** Network representation of all shared guide (top) and target (bottom) contacts.

**Supplementary Fig. 4.**
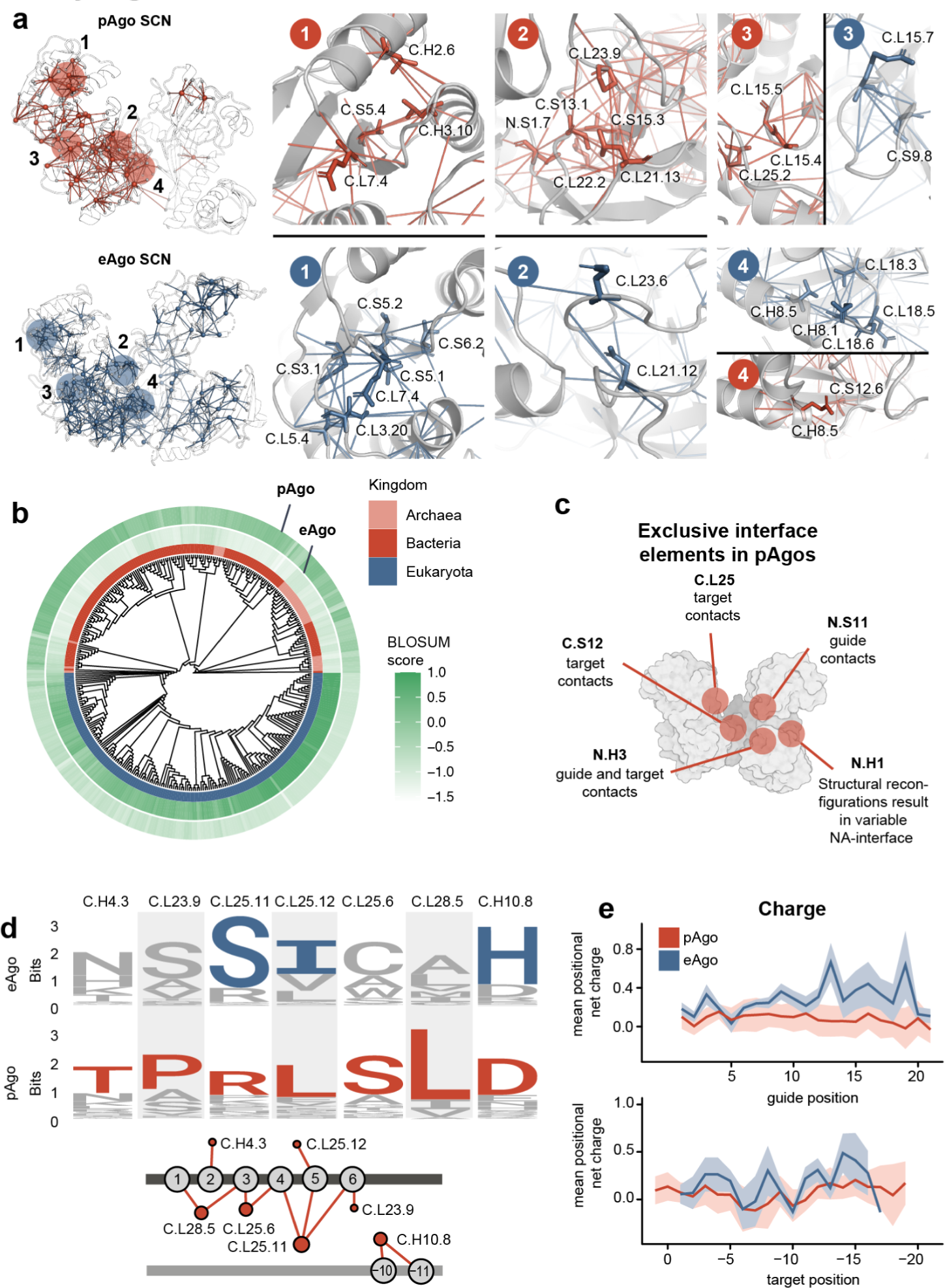
**a** Structural overlay of the pAgo and eAgo SCNs with numbered and highlighted regions in red (TtAgo, PDB: 3DLB) and blue (hAGO2, PDB: 4OLB), respectively. In the MID domain, pAgos show additional distinctly conserved contacts of signature residues in SSEs that form the 5’ binding pocket (C.H2, C.H3), while a cluster of eAgo signature residues can be found in the connecting loops nearby (C.L5, C.L3, C.L7, panel 1). A similar cluster can be found in pAgos mediating contacts underneath the C.L23 loop, which directly contacts the guide in most Argonautes and only carries a few eAgo signature positions in the same region (panel 2). The structural reorientation of the loop C.L15 represents another conserved change from pAgos to eAgos and may imply that these features emerged to enable a new function in eAgos (panel 4). Features like the eAgo cs7 insertion element are also represented in our analysis (panel 5), which lead to a tugged-in conformation of the C.L18 loop, while the same loop is free to engage with the guide or target in pAgos. **b** Normalized BLOSUM score for eAgo and pAgo signature positions against all proteins included in this study with values mapped onto the phylogenetic tree as a heatmap. **c** Examples of exclusive NA interacting features found in pAgos. All NA contact data can be found in Supplementary Data 5. **d** pAgo-specific signatures of the guide interface illustrated in the logo-plot and NA contact network representation. **e** Mean positional net charge of all positions resolved by guide/target position.

**Supplementary Fig. 5.**
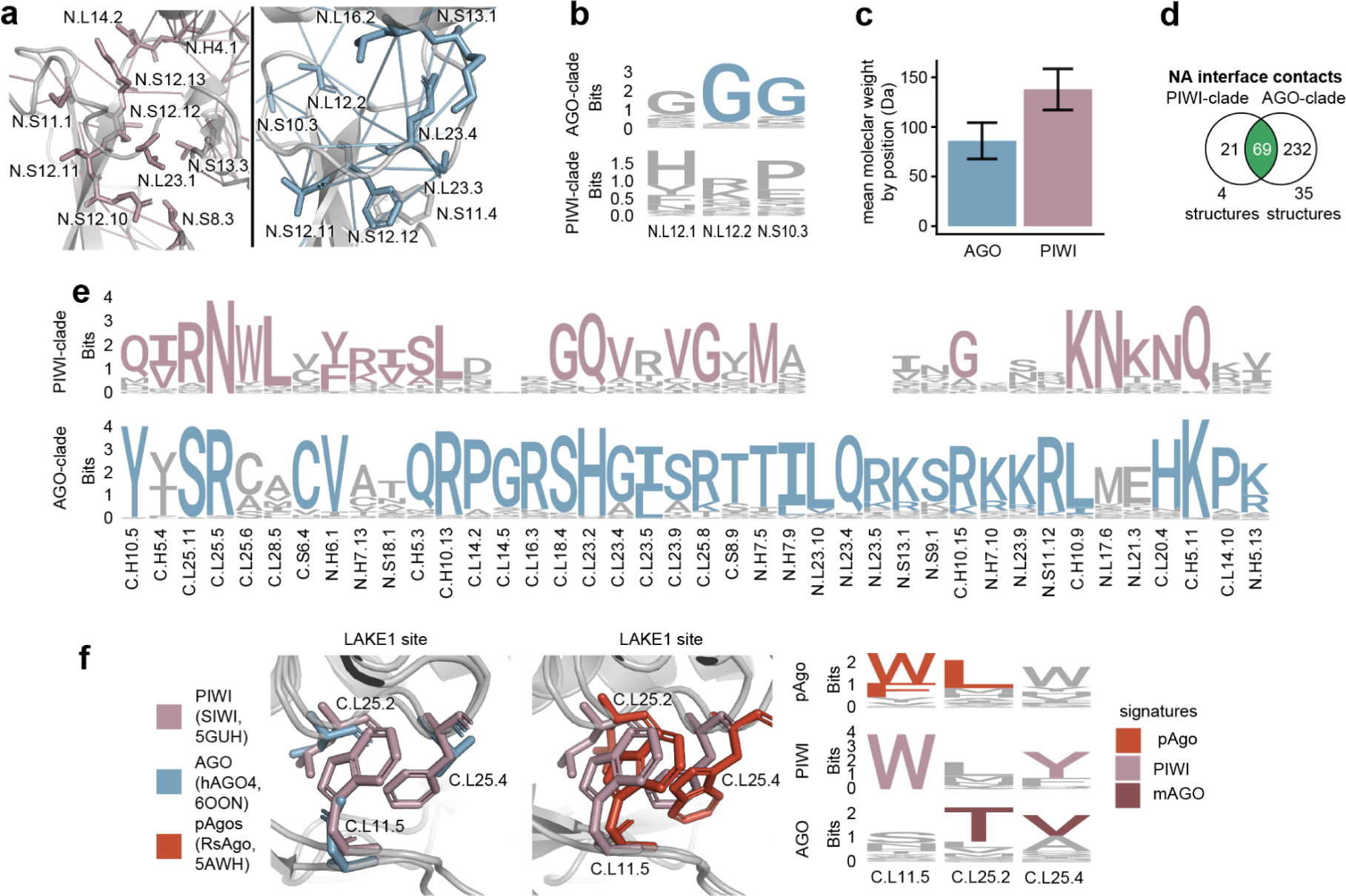
**a** The interface between the PAZ domain and the L2 stalk is mediated by distinct signatures in the AGO and PIWI-clade. The straight PAZ orientation of the AGO-clade is supported by an extended N.L23 interface (e.g. N.L23.4) and the glycine-rich loop N.L12. In AGO, N.S12.12 stabilizes the stalk twist by aromatic interactions with the universal residue N.S11.4. However, in PIWI the tilted PAZ conformation is stabilized by signature positions at the end of the N.S12 strand and N.L14 loop. **b** The N.L12 loop carries bulky residues that further preclude the straight PAZ conformation. Comparison of N.L12 loop residues between AGO and PIWI clade. The positions in the loop are conserved as glycine in the AGO-clade. In PIWI, the loop is not conserved and instead is composed of bulky residues. **c** Mean molecular weight of the positions in **b** for PIWI and AGO. **d** Total NA interface contacts from the available structures in the PDB separated by clade (see Supplementary Data 5 for full annotation of each contact) **e** Sequence logo plot of all PIWI and AGO signature NA-contact positions. **f** Comparison of LAKE1 region in a PIWI, AGO and pAgo representative together with sequence logo plot highlighting signature positions.

**Supplementary Fig. 6.**
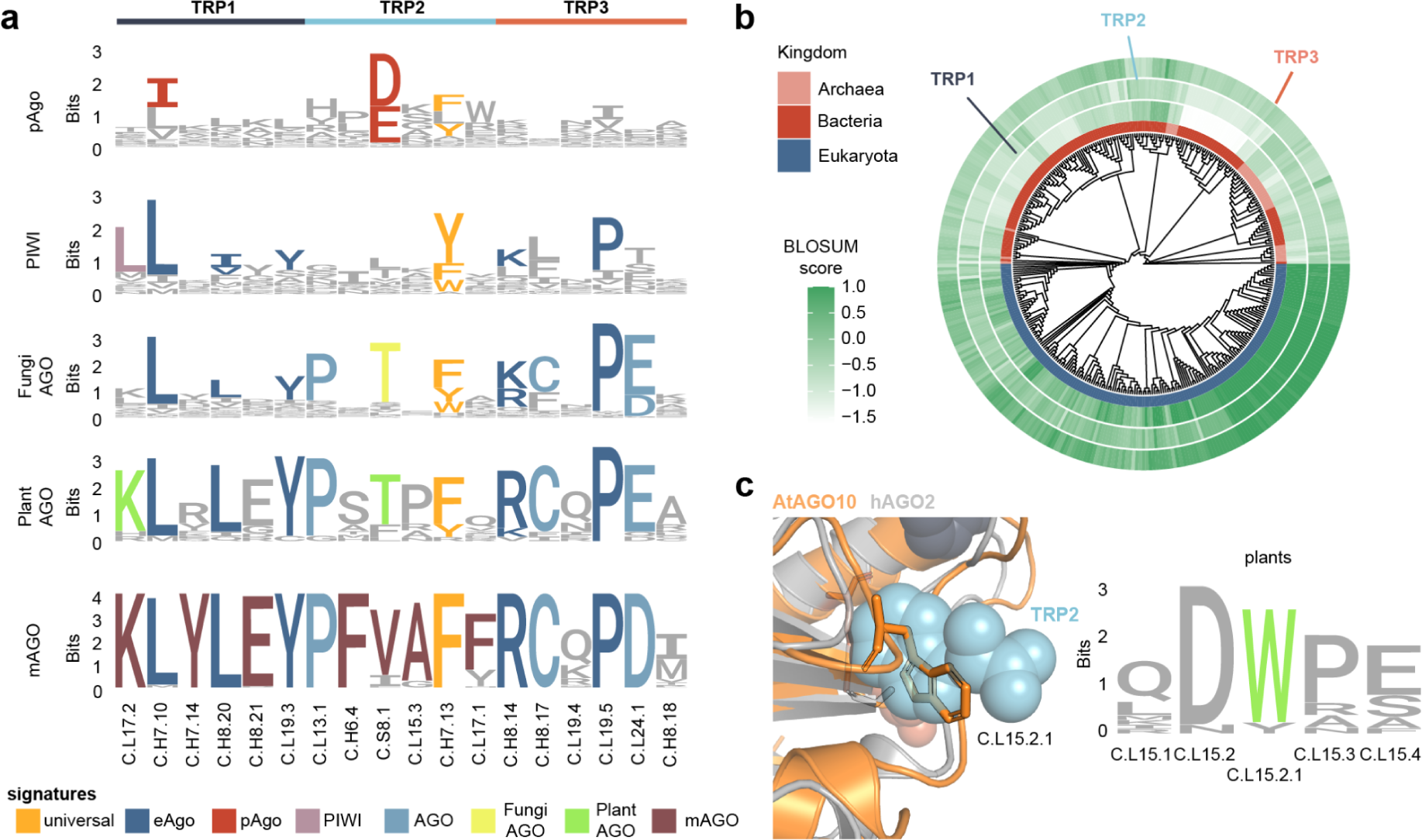
**a** Logo plot of the Argonaute GW-interface positions for multiple distinct phylogenetic groups annotated with signature positions. **b** Normalized BLOSUM score for mAGO consensus GW-interface positions against all proteins included in this study. **c** Structural overlay of AtAGO10 (orange, PDB:7SWF) and hAGO2 (grey, PDB:6CBD) at the TRP2 binding pocket. The C.L15.2.1 (732) position is localized in the GW pocket in AtAGO10. The logo plot of C.L15.2.1 and surrounding positions shows it to be highly conserved as tryptophan in plants.

**Supplementary Fig. 7.**
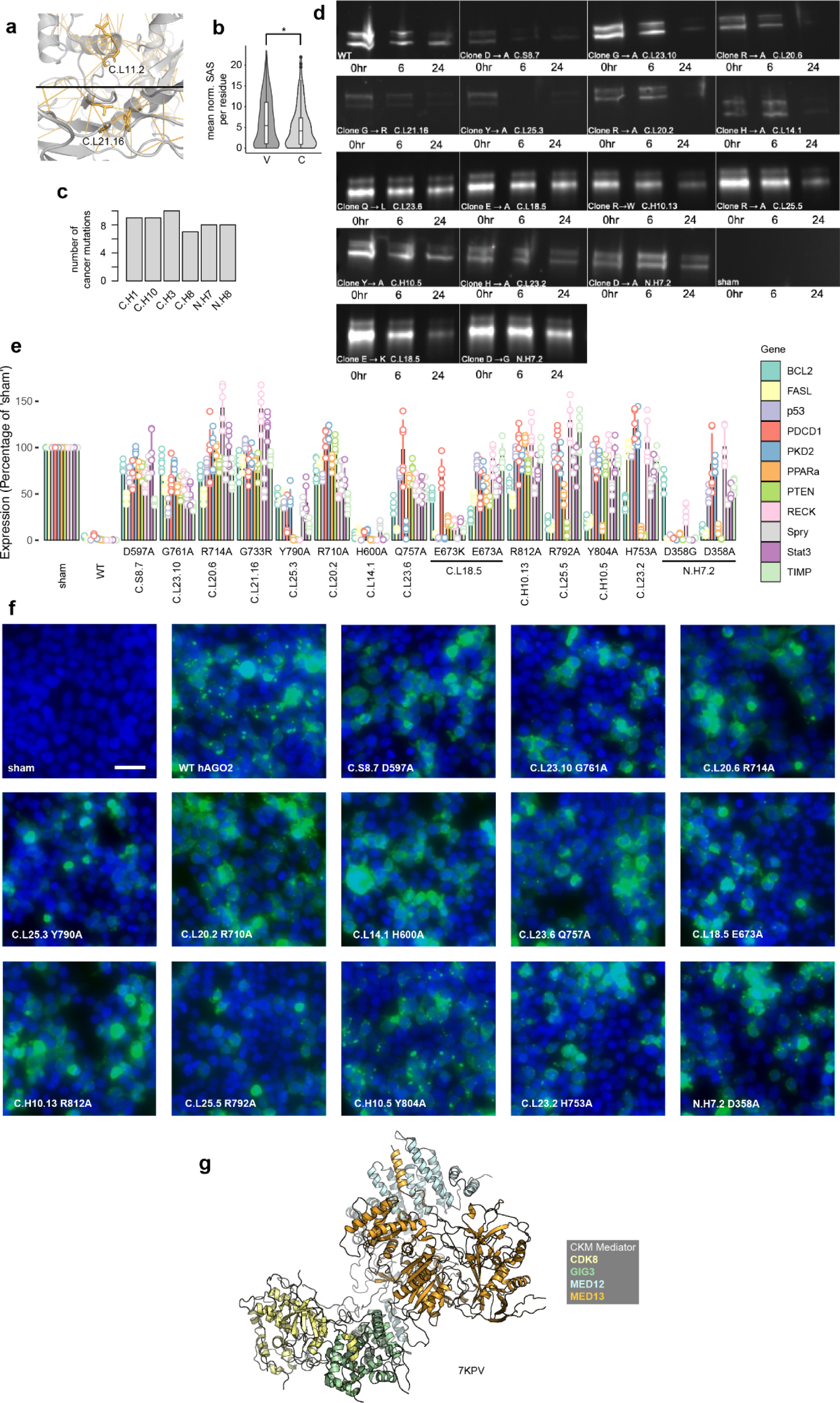
**a** Structural position within the universal SCN of two LESKRES glycine missense mutations. **b** Mean solvent accessible surface (SAS) for variant (V) and cancer (C) mutations. **c** Number of cancer mutations across the six most mutated SSEs **d** Western blotting of CHX-assay for AGO2 stability in hAGO2-GFP mutants. **e** RT-QPCR analysis of relative miR-21 target genes abundance following mutant hAGO2/miR-21 co-transfection. Mutant data from Fig. 6i is repeatedly shown to facilitate comparison. **f** Live cell imaging of hAGO2-GFP (green) containing stress granule (small speckles) formation following Arsenite exposure. Nuclear signal (blue) overlain. Scale bar 20µm. **g** Structure of yeast CKM Mediator complex with annotated subcomponents (PDB: 7KPV).

